# Regulation of sedimentation rate shapes the evolution of multicellularity in a unicellular relative of animals

**DOI:** 10.1101/2021.07.23.453070

**Authors:** Omaya Dudin, Sébastien Wielgoss, Aaron M. New, Iñaki Ruiz-Trillo

**Affiliations:** Institut de Biologia Evolutiva (CSIC-Universitat Pompeu Fabra), Passeig 12 Marítim de la Barceloneta 37-49, 08003 Barcelona, Catalonia, Spain; Institute of Integrative Biology, Department of Environmental Systems Science, ETH Zürich, Zürich, Switzerland; Centre for Genomic Regulation (CRG), The Barcelona Institute for Science and Technology, Dr. Aiguader 88, 08003, Barcelona, Spain; Departament de Genètica, Microbiologia i Estadística, Universitat de Barcelona, Av. Diagonal, 645, 08028 Barcelona, Catalonia, Spain; ICREA, Passeig Lluís Companys 23, 08010, Barcelona, Catalonia, Spain

## Abstract

Significant increases in sedimentation rate accompany the evolution of multicellularity. These increases should lead to rapid changes in ecological distribution, thereby affecting the costs and benefits of multicellularity and its likelihood to evolve. However, how genetic and cellular traits control this process, their likelihood of emergence over evolutionary timescales, and the variation in these traits as multicellularity evolves, are still poorly understood. Here, using isolates of the ichthyosporean genus *Sphaeroforma* - close unicellular relatives of animals with brief transient multicellular life stages - we demonstrate that sedimentation rate is a highly variable and evolvable trait affected by at least two distinct physical mechanisms. First, we find extensive (>300x) variation in sedimentation rates for different *Sphaeroforma* species, mainly driven by size and density during the unicellular-to-multicellular life cycle transition. Second, using experimental evolution with sedimentation rate as a focal trait, we readily obtained, for the first time, fast settling and multicellular *S. arctica* isolates. Quantitative microscopy showed that increased sedimentation rates most often arose by incomplete cellular separation after cell division, leading to clonal “clumping” multicellular variants with increased size and density. Strikingly, density increases also arose by an acceleration of the nuclear doubling time relative to cell size. Similar size- and density-affecting phenotypes were observed in four additional species from the *Sphaeroforma* genus, suggesting variation in these traits might be widespread in the marine habitat. By resequencing evolved isolates to high genomic coverage, we identified mutations in regulators of cytokinesis, plasma membrane remodelling, and chromatin condensation that may contribute to both clump formation and the increase in the nuclear number-to-volume ratio. Taken together, this study illustrates how extensive cellular control of density and size drive sedimentation rate variation, likely shaping the onset and further evolution of multicellularity.

## Introduction

The emergence of multicellularity from single-celled life represents a major transition which has occurred many times independently across the tree of life (Grosberg & Strathmann, 2007; Knoll, 2011; Leigh et al., 1995; Niklas & Newman, 2013; Parfrey & Lahr, 2013; Rokas, 2008; Ruiz-Trillo et al., 2007; Sebé-Pedrós et al., 2017). Multicellularity can arise either by aggregation of single cells that come together, or from single cells that are maintained together clonally after division (Bonner, 1998; Tarnita et al., 2013; Wielgoss et al., 2019). The unicellular and intermediate multicellular ancestors which led to present-day multicellular organisms have long been extinct (Grosberg & Strathmann, 2007), obscuring direct investigation of how multicellular life has emerged. However, several strategies have been used to study the emergence of multicellularity, including the use of experimental evolution (EE) approaches and the investigation of novel non-model organisms at pivotal positions in the tree of life.

For example, EE under controlled conditions allows selection for diverse phenotypes (Elena & Lenski, 2003; Kawecki et al., 2012), including multicellularity (Herron et al., 2019; Koschwanez et al., 2013; Ratcliff et al., 2012, 2013, 2015). Using EE, Ratcliff and colleagues repeatedly observed the evolution of a simple form of multicellularity in *Saccharomyces cerevisiae* and *Chlamydomonas reinhardtii* in response to gravitational selection (Oud et al., 2013; Ratcliff et al., 2012, 2013, 2015). Similarly, multicellularity emerged in yeast as a mechanism to improve the use of public goods (Koschwanez et al., 2013). In all these cases, cells form clumps by incomplete separation of daughters from mother cells, instead of by post-mitotic aggregation (Fisher et al., 2013; Queller et al., 2003; Strassmann et al., 2011).

Alternatively, non-model organisms with key evolutionary positions can be used to better understand the emergence of multicellularity. In particular, the study of unicellular holozoans (Figure 1A), the closest unicellular relatives of animals, revealed that these organisms contain a rich repertoire of genes required for cell adhesion, cell signalling and transcriptional regulation, and that each unicellular holozoan lineage uses a distinct developmental mode that includes transient multicellular forms (Brunet et al., 2019; Brunet & King, 2017; Parra-Acero et al., 2020; Pérez-Posada et al., 2020; Ruiz-Trillo & de Mendoza, 2020; Sebé-Pedrós et al., 2017). For instance, the choanoflagellate *Salpingoeca rosetta* can form clonal multicellular colonies through serial cell division in response to a sulphonolipid of bacterial origin (Alegado et al., 2012; Fairclough et al., 2010; Levin et al., 2014), whereas the filasterean *Capsaspora owczarzaki* can form multicellular structures by aggregation (Sebé-Pedrós et al., 2013). Ichthyosporeans display a coenocytic life cycle unique among unicellular holozoan lineages and pass through a short and transient clonal multicellular life-stage prior to the release of new-born cells (A. de Mendoza et al., 2015; Dudin et al., 2019; Glockling et al., 2013; L. Mendoza et al., 2002; Ondracka et al., 2018). Despite their pivotal phylogenetic position, their rich “animal” genetic toolkit and the capacity to undergo transient multicellularity, to date, unicellular holozoans lineages and EE, have never been combined.

**Figure 1.**
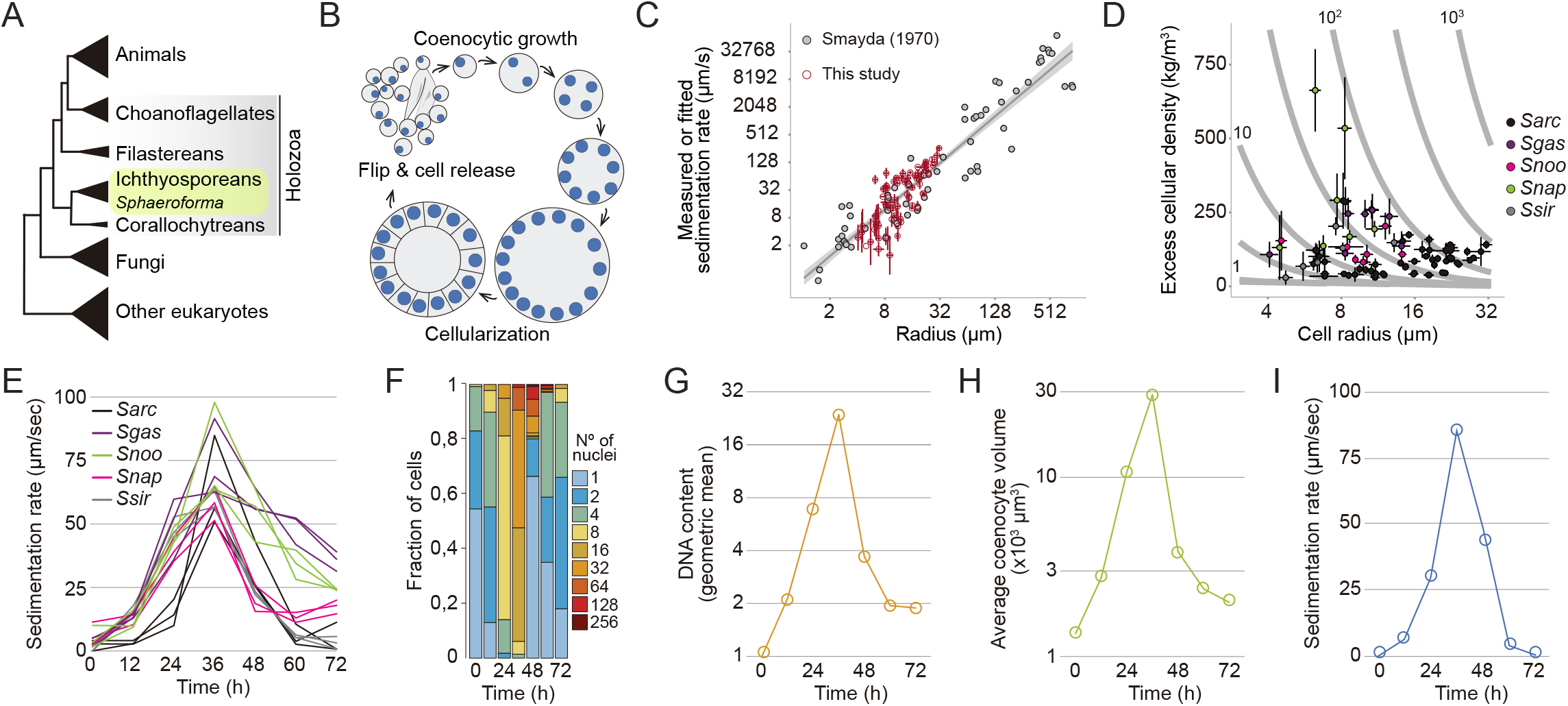
Sedimentation dynamics of *Sphaeroforma arctica* coenocytes during the life cycle. (A) Cladogram representing the position of ichthyosporeans including *Sphaeroforma* species within the eukaryotic tree. (B) Schematic representation of the coenocytic life cycle of *S. arctica*. (C) Data derived from Figure 1 (SMAYDA & J., 1970) (grey points) were used to scale velocity measurements determined in our sedimentation assay to physical (µm/s) units (red points) (Methods). Error bars represent the 95% CI for each unique genotype, timepoint, and temperature measurement presented in our study (N=3 for each of 1 or 2 independent replications). Values were log-transformed prior to calculation of error. This figure is to illustrate where our data fit in the scheme of known plankton sedimentation rates. For our best estimations of cellular density and velocity in meters per second, a subset of this data from Smayda’s Appendix Table 1 was used (Methods). (D) Relying on the Smayda dataset as reference, we used measurements of sedimentation rate from our assay along with cellular perimeter measurements to calculate maximum likelihood estimates of excess cellular density. See Methods and Figure 1 – Source Data. These estimates are plotted on a landscape illustrating the relationship between density and size on sedimentation rate (grey contour lines). Grey contour lines represent the predicted settling velocity of each genotype in pure water (excess density = 1000 kg/m^3^) in units µm/s. (E) Sedimentation rates of *Sphaeroforma* during the life cycle at 17°C. Every trace represents an independent experiment. (F) Distributions of nuclear content of *S. arctica* cells during the life cycle at 17°C, measured by microscopy (n > 500 cells per timepoint). (G) Quantification of mean DNA content per time point (measured by geometric mean) for cells grown in marine broth at 17°C (n > 500 cells per timepoint). (H) Average coenocyte volume per timepoint at 17°C (n = 100 coenocytes per timepoint).(I) Sedimentation rates of *S. arctica* coenocytes during the life cycle at 17°C.

Beside the formation of new biological structures and increases in organismal size, the emergence of multicellularity is frequently accompanied with an increase in sedimentation rate. Indeed, *S. cerevisiae* snowflake yeast, multicellular *C. reinhardtii* and *S. rosetta* colonies sediment faster when compared to their unicellular counterparts (Ratcliff et al., 2012, 2013; Thibaut Brunet – Personal communication). Such correlation has been described in several marine phytoplankton species where multicellular life-stages show faster-sedimentation than unicellular ones (Beardall et al., 2009; Eppley et al., 1967; Finkel et al., 2010; Smayda, 1971). This phenotype has a large impact on where in the water column these microbes proliferate (Beardall et al., 2009; Friebele et al., 1978; Gemmell et al., 2016; N. B. Marshall, 1954), and thus is presumed to be under strong genetic control and selective pressure. Despite its capacity to affect the depth at which marine species flourish, the role of sedimentation rate, the potential impact of its variation and its connection to the emergence of multicellularity has not been systematically analysed across unicellular marine organisms, including in the closest unicellular relatives of animals with transient multicellular life-stages. Here, using EE, we characterize how regulation of sedimentation rate can influence the emergence of stable multicellular life-forms in the ichthyosporean *Sphaeroforma* genus, a close unicellular relative of animals.

## Results

### *Sphaeroforma* species exhibit large variation in sedimentation rates

Similar to other ichthyosporeans, *Sphaeroforma* species proliferate through continuous rounds of nuclear divisions without cytokinesis to form a multinucleated coenocyte (Glockling et al., 2013; W. L. Marshall et al., 2008; Ondracka et al., 2018; Suga & Ruiz-Trillo, 2013). *Sphaeroforma* coenocytes then undergo a coordinated cellularization process leading to the formation of a transient multicellular life-stage resembling an epithelium (Dudin et al., 2019). This layer of cells then detaches and cell-walled new-born cells are released to the environment (Figure 1B) (Dudin et al., 2019). The entire life cycle prior to cellularization occurs in highly spherical (multinucleated) cells.

Extensive literature has documented a positive correlation between cell size and sedimentation rate, including during the life cycle of marine phytoplankton (Figure 1C) (SMAYDA & J., 1970; Smayda, 1971).This is consistent with Stoke’s law, which shows that the relationship between a spherical particle’s terminal sedimentation rate ***v*** in a fluid and its radius *R* should be determined by

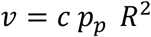

Where ***c*** represents the scaled ratio of gravitational to viscosity constants, and ***p_p_*** is the difference between the particle’s and the fluid’s densities (see Methods). Therefore, even small shifts in a particle’s radius will lead to pronounced (i.e., quadratic) changes in sedimentation rate. Similarly, for particles sitting near the buoyancy threshold, small changes in density can lead to proportionally large changes in settling rate (Figure 1D).

Due to the nature of the coenocytic life cycle (Figure 1B), which is associated with an increase in the number of nuclei and coenocyte volume, we expected to observe an increase in cellular sedimentation rates over time (Allen, 1932; N. B. Marshall, 1954; Waite et al., 1997).To better understand this relationship, throughout this study, we conducted overlapping experiments characterizing cell volume and sedimentation rates of *Sphaeroforma* species for cultures growing at 17°C for 72 hours. For certain replicates of this core dataset, we included measurements of various genetic variants, the temperature dependence of phenotypes, fitness, as well as the speed of nuclear duplication. Overall, the measurements we report are highly reproducible with >95% variance explained for replicate measurements across phenotypes (Figure 1 – figure supplement 1A and Figure 1- data source 1 – 4) (see Methods). This reflected a high heritability of the different phenotypes in a given environment.

To begin, we measured the sedimentation rate of five different *Sphaeroforma* species that have been isolated from different habitats either free-living or host-derived, namely *S. arctica*, *S. sirkka*, *S. napiecek*, *S. gastrica* and *S. nootakensis*, using a quantitative sedimentation rate assay based on changes in optical density over time (Ducluzeau et al., 2018; Dudin et al., 2019; Hassett et al., 2015; Jøstensen et al., 2002; W. L. Marshall et al., 2008; W. L. Marshall & Berbee, 2013). According to these estimates, cell sedimentation rates varied greatly, from between 0.4 to 125 µm per second (i.e., up to 0.45 meters per day) (Figure 1E). This broad variability over the life cycle and among *Sphaeroforma* species suggested that appreciable changes in size and/or cellular density should accompany different stages of the life cycle.

### The transient multicellular life stage of *S. arctica* is associated with an increase in sedimentation rate

To better understand the cellular basis of sedimentation rate variation, we focused on *S. arctica,* the most studied *Sphaeroforma* species to date (Dudin et al., 2019; Jøstensen et al., 2002; Ondracka et al., 2018). Using fixed-cell imaging, we observed that synchronized cultures of *S. arctica* undergo a complete life cycle in about 48 hours and can reach up to 256 nuclei per coenocyte before undergoing cellularization and releasing new-born cells (Figure 1F). Prior to this release, all cellular division occurs in highly spherical mother coenocytes. Consistent with previous results (Ondracka et al., 2018), nuclear division cycles were periodic during coenocytic growth and occurred on average every 9-10 hours, as measured indirectly from changes in DNA content (Figure 1G). Average cell volume increased throughout the coenocytic cycle to reach its maximum value at 36 hours prior to the release of new-born cells (Figure 1H). Similarly, the sedimentation rate increased up to >300-fold the initial value after 36 hours (Figure 1I). Altogether, we observe that every cycle of nuclear division is associated with a significant increase in nuclear content, volume and sedimentation rate with a distinct peak just prior to cell release. As the transient multicellular life-stage of *S. arctica* occurs during the latest stage of cellularization and ahead of cell-release (Dudin et al., 2019), our results suggest that it is tightly associated with an increased sedimentation rate.

### Experimental evolution of fast-settling mutants

Given how cell size and division across the cell cycle are regulated, we reasoned that these variable traits should also be heritable and hence evolvable. To test this, we conducted an evolution experiment to generate mutants with increased sedimentation rates (Herron et al., 2019; Ratcliff et al., 2012, 2013). Briefly, 10 independent cultures of *S. arctica* (S1 to S10) originating from the same ancestral clone (AN) (Figure 2A), were diluted in fresh marine broth medium at 17°C (Figure 1C). Selection was performed every 24 hours by allowing the cultures to sediment in tubes for 2 minutes before transferring and propagating the fastest-settling cells in fresh medium (Figure 2A). Experimental evolution was continued for 56 transfers (8 weeks) or about 364 nuclear generations and a frozen stock was conserved every 7 transfers (1 week) (see Methods) (Figure 2—figure supplement 1A, Figure 2- Source data 1). Most lineages exhibited a distinct clumping phenotype to varying degrees across all evolved populations (Figure 2B). This phenotype has been never observed in any *Sphaeroforma* culture, despite variation of culturing conditions across labs, and the hundreds of routine passages without selection, thus suggesting that the appearance of such phenotype is linked to our selection regime. The clumps of cells were not maintained by ionic forces or protein-dependent interactions, however could be separated by mild sonication without leading to cell lysis (Figure 2- figure supplement 1B). To assess when sedimentation rates increased during the selection process for each evolved population, we synchronized cultures using mild sonication before dilution in fresh medium and allowed them to undergo a complete life cycle before measuring the sedimentation rate (Figure 2C). We observed that populations S1, S4, and S9 had the highest sedimentation rates at the end of the evolution experiment (Figure 2C). We observed a dramatic increase in sedimentation rate already after 14 transfers for population S1 (∼91 generations) (Figure 2C). To assess and compare the variability in sedimentation rates among evolved lineages, we isolated and characterized a single clone from each evolved culture. Our results show that evolved clones all settled significantly faster than the common ancestor, but that there was stark variation in sedimentation rates at the end of the EE (Figure 2D). Isolates from lineages S1, S4 and S9 settled the fastest upon sedimentation (Video 1), while clones from lineages S2, S3, S5 and S7 were intermediate, and lineages S6, S8 and S10 settled the slowest (Figure 2D). Taken together, our results show that we can rapidly and routinely evolve fast-settling mutants in *S. arctica* using experimental evolution, but that the outcome is not uniform across lineages.

**Figure 2.**
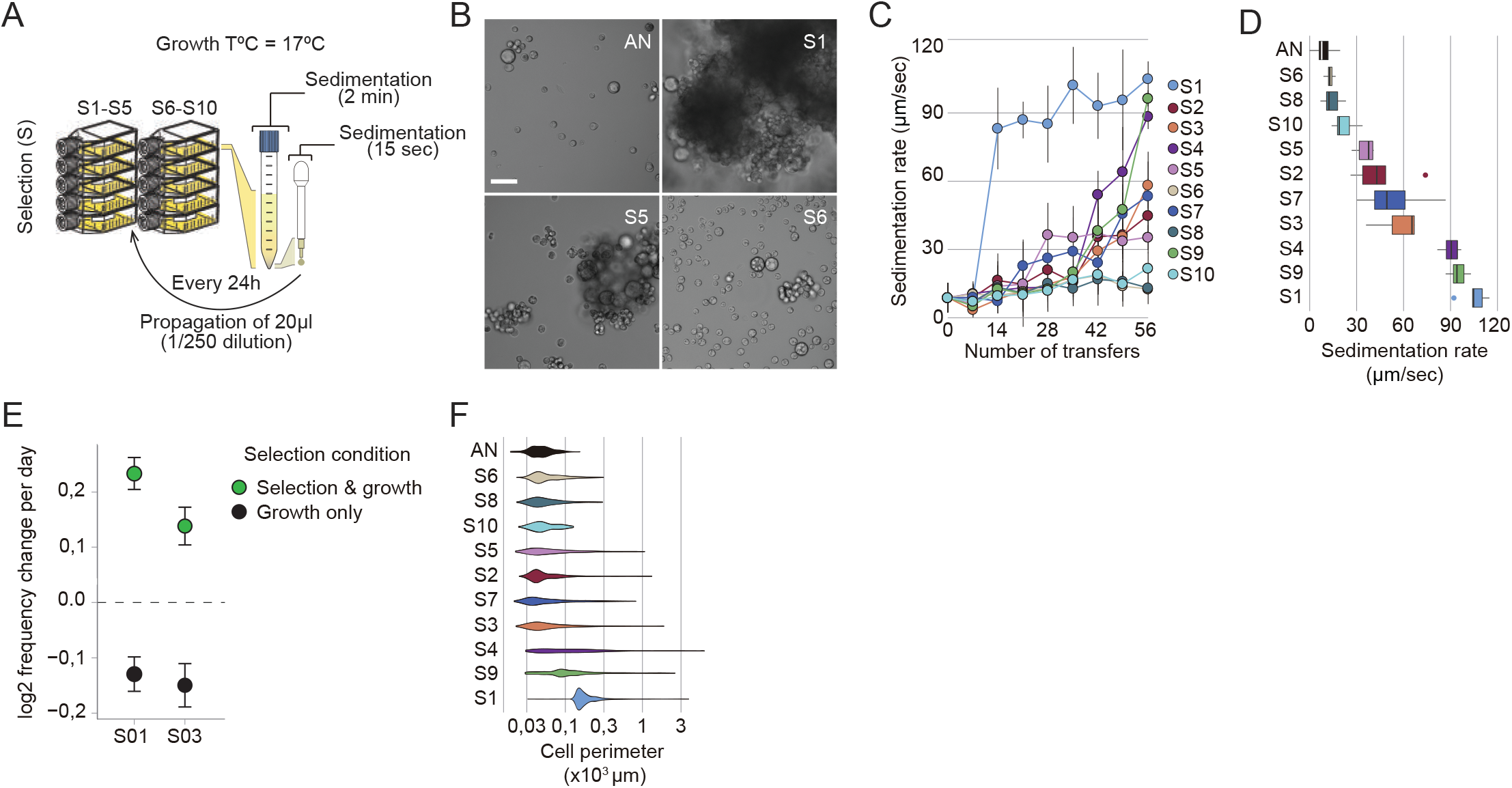
Rapid evolution of fast-settling and clumpy mutants. (A) Schematic representation of the experimental evolution design using sedimentation as a selective pressure. All 10 replicate populations (S1-S10) were isolated from the same ancestral culture (AN) and were subject to 56 transfers over 8 weeks of selection. (B) Representative images of the ancestral culture (AN) and three evolved mutants (S1, S5, S6) with varying clumping capability after 5 days of growth. Bar, 50µm. (C) Sedimentation rates of *S. arctica* evolved populations per number of transfers shows rapid emergence of fast-settling phenotypes, particularly in S1. (D) Sedimentation rates of single clones of *S. arctica* evolved mutants after 56 rounds of selection displays vast variations in sedimentation rate. Associated with Video 1. (E) Head-to-head competitions between the clumpy evolved isolates S01 & S03 with the ancestral strain (AN) show that sedimentation rate is adaptive in the conditions of our evolution experiment. The rate of change of the frequency of the clumpy phenotype per day in log two units was determined by least-squares regression. Error bars represent the 95% confidence interval based on 500 calculations of this statistic obtained by resamplings of the observed binomal count frequencies for each data point. (F) Clump size distribution of evolved clones shows that fast-settling mutants (S1, S4, S9) show bigger clump size (n > 800 measurements per timepoint).

To assess whether fast sedimentation is adaptive in our EE set-up, we independently competed two fast sedimenting isolates, S01 and S03, against their common ancestor (AN). For this relative fitness assay, we mixed mono-cultures of either S01 or S03 with AN at a ∼1:1 ratio, and subjected them to the same selective regime as during EE. Parallel cultures of each 1:1 mix grown and maintained without sedimentation selection served as experimental control (see methods). Our results show that the proportion of clumpy S01 and S03 cells both significantly increased over time (a log2 selection coefficient of 0.23 and 0.13 per day for S01 and S03, respectively, p<1e-7), suggesting this trait was adaptive in our experimental set-up (Figure 2E, Figure 2- figure supplement 1D, Figure 1 – figure supplement 1A, Figure 1- data source 4). In contrast, S01 and S03 cells decreased in frequency relative to the AN strain in the control growth experiment without sedimentation (a log2 selection coefficient of -0.13 and -0.15 per day for S01 and S03, respectively, p<1e-7), suggesting that the heritable changes which these strains bore were deleterious in normal laboratory propagation conditions (Figure 2E, Figure 2- figure supplement 1D).

We next sought to characterize the cellular mechanisms giving rise to variation in cellular sedimentation rates in evolved clones. Using quantitative microscopy, we show that clump perimeter and average number of cells observed per clump correlated highly with sedimentation rates (Figure 2E and Figure 2- supplement figure 1C, D and E). Indeed, across all experiments reported in this study, we found that using Stokes’ law, a single globally fitted density parameter and our size measurements, could explain the majority of variance in sedimentation rate (R^2^ = 0.69, RMSE (as a proportion of the range in observations) = 18%).

### Fast-sedimenting mutants form clonal clumps

Earlier, we defined three distinct developmental stages of *S. arctica* life cycle: (i) cell growth with an increase in coenocyte volume, (ii) cellularization which coincides with actomyosin network formation and plasma membrane invaginations, and (iii) release of new-born cells (Figure 1B) (Dudin et al., 2019). A key developmental movement named “flip”, defined by an abrupt internal morphological change in the coenocyte, can be used as a reference point to characterize life cycle stages. Prior to this event (pre-flip), actomyosin-dependent plasma membrane invaginations occur, while afterwards (post-flip) the cell wall is formed around individual cells prior to their release from the coenocyte (Dudin et al., 2019). Using time-lapse microscopy at 12°C, we observed that clump formation in fast-settling mutants coincides with cell release (Figure 3A, Video 2).

**Figure 3.**
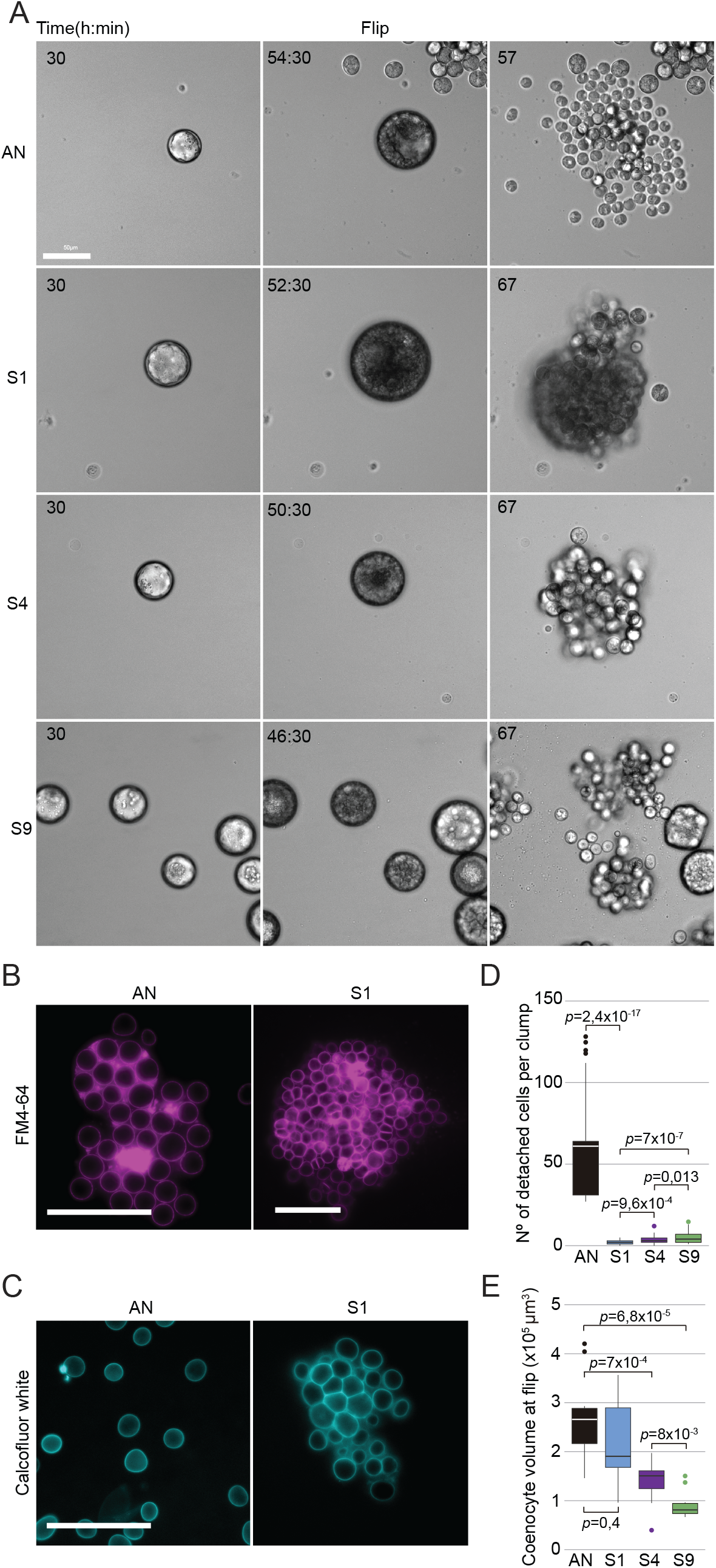
Clumps are formed by incomplete cell-cell separation in fast-settling mutants. (A) Time-lapse images of the life cycle of *S. arctica* ancestral strain (AN) and the fast-settling mutants (S1, S4, S9) at 12°C, show that clumps are observed concomitantly with release of new-born cells. Associated with Video 2. Bar, 50 µm (B) Plasma membrane staining using FM4-64 show that plasma membrane invaginations during cellularization seems to occur normally in fast-settling mutants. Associated with Video 3. Bar, 50µm. (C) Cell-wall staining using calcofluor-white indicate cells are still separated by a cell-wall inside fast-settling clumps, suggesting that clumps are formed post-flip. Bar, 50µm. (D) Number of cells detaching per clumps at cell release and for the following three hours, measured from time-lapse movies (n = 50 coenocytes at cell-release per strain). (E) Coenocyte volume at flip, measured from time-lapse movies, show that S4 and S9 are significantly smaller when compared to AN and S1 (n = 50 coenocytes at flip per strain).

Importantly, no new cell aggregation processes were detected after cell release, suggesting that all clumps formed either prior to or concomitantly with the release of new-born cells (Video 2). To examine whether clumps formed due to defects at the level of either plasma membrane or cell wall, we stained plasma membranes using FM4-64 and cell walls using calcofluor-white. We found that the process of plasma membrane invaginations during cellularization appears to be unchanged prior to flip (Figure 3B- Video 3), and that all new-born cells, even in the clumps, have a distinctive cell wall surrounding them (Figure 3C, Figure 3- figure supplement 1A). These results show that clumps are formed post-flip in fast-settling mutants.

As no cells appear to aggregate after release of new-born cells, our results suggest that clumps are maintained together in a clonal form. To further confirm this, we sonicated both the ancestor (AN) and S1 clumps and stained them separately with two distinct fluorescent dyes prior to mixing them in a 1:1ratio (Figure 3- figure supplement 1B). After one complete life-cycle, we observed that out of 198 clumps, the largest 183 contained only evolved S1 derivates, whereas ancestral cells only randomly, and sporadically formed clumps under the conditions of the experiment (a total of 12 small clumps). Only three of the clumps (∼1.5%) contained cells of both colours (Figure 3- figure supplement 1B), and each only contained a single AN cell trapped inside a smaller S1 clump. In a control experiment with two differentially stained S1 cultures, we observed an almost identically small number (i.e., ∼1.7%) of mixed clumps (Figure 3- figure supplement 1B), hence, it appears that mixes of cells happen only sporadically at low frequencies by random association, irrespective of genotype. Altogether, our results suggest that the evolved clump phenotype is not a result of spontaneous cell aggregation, and instead arises from incomplete detachment between cells.

We next examined the kinetics of cellularization and the process by which cells propagate as part of clumps. We counted the number of cells detaching from clumps at cell release and observed variable detachment across all three fast-settling mutants. The clumpiest isolate, population S1, was the least prone to cell detachment (Figure 3A and D, Videos 2, 3). Intriguingly, despite exhibiting similar sedimentation rates, S4 and S9 clumps showed a higher detachment frequency than S1 (Figure 3A and D). Image analysis of time-lapse movies also showed that life-stage durations varied among fast-settling mutants at 12°C, with S1, S4, and S9 initiating cellularization and undergoing flip, respectively, 5.5 hours, 7.5 and 10.5 hours earlier than the ancestor (Figure 3- figure supplement 1C-D). Post-flip duration also varied significantly among mutants, with S1 and S9 requiring more time to release cells compared to S4 (Figure 3- figure supplement 1C-D). While many aspects of the replication cycle dynamics were variable for these mutants, the duration of cellularization was fairly invariant (∼9 hours) (Figure 3- figure supplement 1C-D). Finally, measurements of coenocyte volume show that S4 and S9 coenocytes undergo flip at substantially smaller volumes (∼1,8 and ∼3,3x times smaller, respectively) compared to AN or S1 (Figure 3E). These results show that, despite their shared capacity to clump and similar sedimentation rates, fast-settling mutants exhibit significant variability in their life cycle dynamics, with S4 and S9 mutants initiating cellularization earlier, dispersing from clumps with a higher frequency, and undergoing flip and cell release at smaller coenocyte volumes compared to the S1 mutant.

### Increased nuclear number-to-volume ratio leads to faster sedimentation

Above we observed that S4 and S9 mutants can sediment as fast as S1 despite their smaller coenocyte volumes, suggesting an alternative regulation mechanism of sedimentation rate. Across all experiments reported in this study, we found that ∼31% of the variance in observations of sedimentation rate could not be explained by cell size only, suggesting that cellular density might also contribute to this variation (Eppley et al., 1967; SMAYDA & J., 1970; Smayda, 1971). From Stoke’s law, we calculated that *excess cellular density*, i.e., cellular density minus that of distilled water (1000 kg/m^3^), might vary between 40 and 300 kg/m^3^ for *S. arctica* wild-type and evolved clones across their life cycle – the upper limits between the densities of pure protein and pure cellulose. Values reached ∼650 kg/m^3^ for wild *S. nootakensis* (*Snoo*) soon after cellularization, approaching the excess density of pure nucleic acid (Figure 1D). During cell cycle stages prior to cellularization, when cells were most spherical (<36 hours), excess density varied from 40-200 kg/m^3^ across *Sphaeroforma* isolates.

To better characterize the relationship between sedimentation rate, cell cycle and size we performed higher resolution measurements of sedimentation rates over the complete life cycle of the ancestor and all three fast-settling mutants at 12°C and 17°C. Consistent with their capacity to form clumps, we observed that the sedimentation rate of all fast-settling mutants increases during growth but, unlike the ancestor, does not recover after cell release to their original levels (Figure 4A and figure 4-figure supplement 1A). Interestingly, we noticed that individual S4 and S9 coenocytes sediment faster (∼2,5x and ∼1,6x respectively) than S1 or AN even before clump-formation (24 to 36 hours timepoints) (Figure 4A and figure 4-figure supplement 1A). Such increase in sedimentation rate was not due to a rise in cell size or change in cell shape as both S4 and S9 exhibit smaller cell perimeters throughout the cell cycle (Figure 4B, C and figure 4-figure supplement 1B, C). Rather, excess cellular density estimations show that both S4 and S9, even prior to cell release and clump formation, tend to be on average 3x denser when compared to the ancestor (Figure 4D and figure 4-figure supplement 1D). Altogether, these results show that both cell size and cell density contribute to sedimentation rate variation in *S. arctica*.

**Figure 4.**
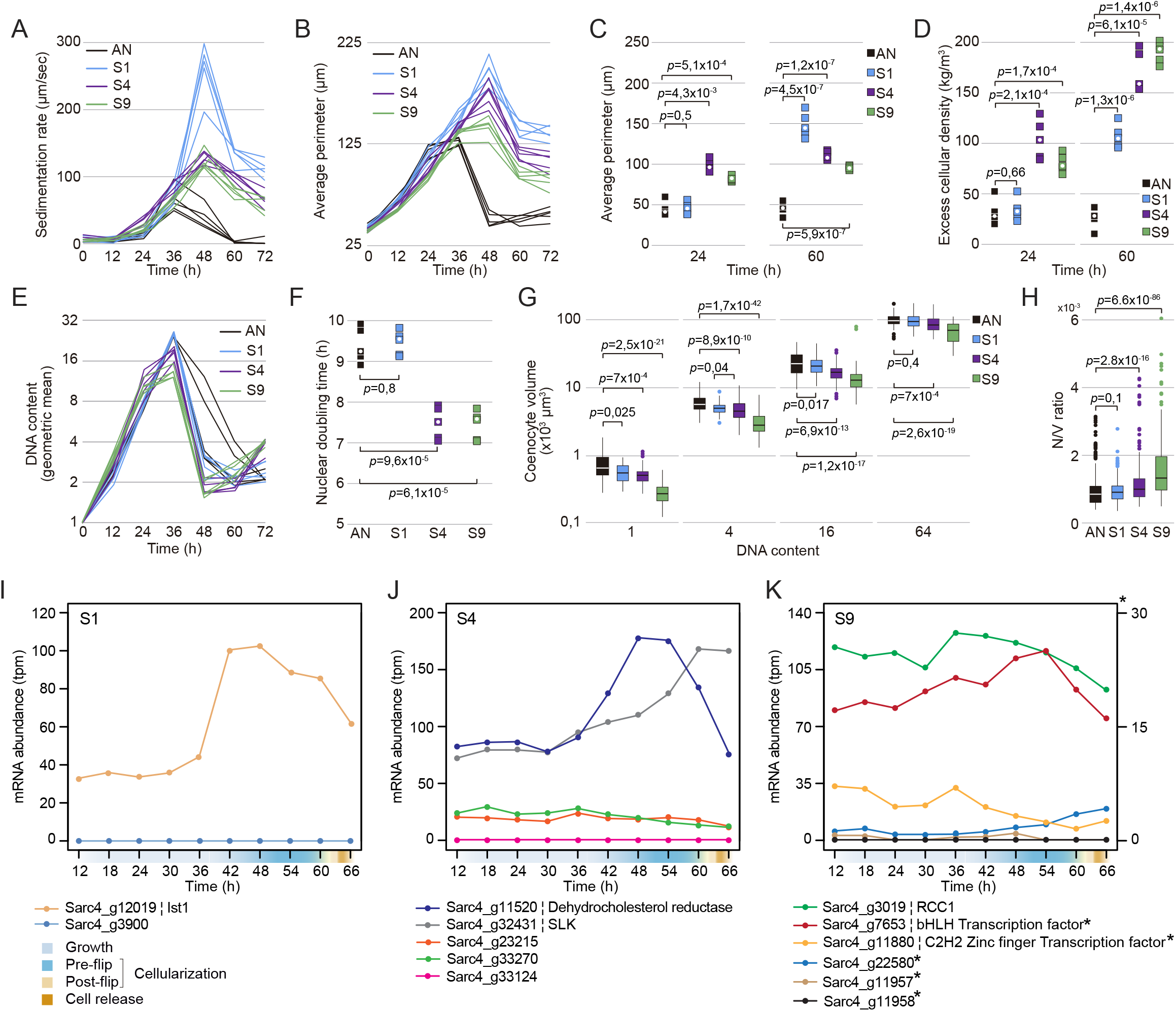
Sedimentation rates variation in fast-settling mutants is associated with variation in cell size and cellular density. (A) Sedimentation rates of *S. arctica* AN and evolved mutants during the life cycle at 17°C. Every trace represents an independent experiment. (B) Average perimeter measured from fixed cells every 12 hours over a complete life cycle of 72 hours at 17°C shows that fast-settling mutants increase their size upon-cell release. Every trace represents an independent experiment (n > 180 measurements per timepoint for each independent experiment). (C) Average perimeter of fast-settling cells and clumps at 24 and 60 hours respectively show that S4 and S9 cells and clumps have a smaller size when compared to S1. Every square represents an independent experiment, and the white circle represents the median (n > 180 coenocytes per timepoint for each independent experiment). (D) Excess cellular density of fast-settling individual coenocytes (before cellularization) and clumps (after cell-release) at 24 hours and 60 hours respectively show that S4 and S9 single cells are denser when compared to S1 and AN. Every square represents an independent experiment, and the white circle represents the median. (E) Quantification of mean DNA content per time point for fast-settling mutants grown in marine broth at 17°C. Every trace represents an independent experiment (n > 400 coenocytes per timepoint for each independent experiment). (F) Nuclear doubling time, calculated by linear regression of mean nuclear content at timepoints from 0 hr to 24 hours. Every square represents an independent experiment, and the white circle represents the median (n > 400 coenocytes per timepoint for each independent experiment). (G) Boxplots of cell volume measurements of DAPI-stained fixed cells. For 1-, 4-, 16-, and 64-nuclei cells. Cells with one nucleus represent new-born cells at the end of the experiment (n > 80 coenocytes per DNA content). (H) Boxplots of nuclear number-to-volume ratio of DAPI stained cells show significant increase for S4 and S9 fast-settling mutants. Every square represents an independent experiment, and the white circle represents the median (n > 600 coenocytes per strain). (I-K) Temporal transcript abundance of genes mutated in fast-settling phenotype across the native life cycle of *S. arctica*.

As cell size and nuclear division cycles are decoupled in *S. arctica* (Ondracka et al., 2018), we reasoned that increased cell density in S4 and S9 could be caused by an acceleration of nuclear divisions leading to a rise in the number of nuclei per volume. Using DAPI staining to label nuclear DNA, we observed that S4 and S9 undergo nuclear duplication faster (∼2 hours) than both AN and S1 (Figure 4E, F and figure 4-figure supplement 1E to H). By carefully examining the volumes of coenocytes containing the same number of nuclei at the single-cell level, we show that for the same nuclear content, S4 and S9 tend to be 30-45% smaller in volume when compared to the ancestor (Figure 4G and figure 4-figure supplement 1I). Consequently, both S4 and S9 exhibited the highest number of nuclei per volume (nuclear number-to-volume ratio) (Figure 4H and figure 4-figure supplement 1J). Taken together, these results argue that cell density can contribute an appreciable amount to cellular sedimentation rates (up to ∼50 µm/s), and that mechanistically this could arise by faster nuclear doubling times relative to cell size.

### Evolved genetic variation correlating with fast sedimentation

Up to now, our results suggest that *S. arctica* mutants evolved faster sedimentation using two strategies: (i) clump formation, and (ii) increased nuclear number-to-volume ratio. We found that sedimentation rate variation was highly heritable, persisting for 780 generations of passaging for all 10 isolates without selection for sedimentation phenotype, suggesting that the phenotypes have a genetic basis. To test this, we resequenced the whole genomes of both the ancestral clone (AN) and one evolved clone per lineage (S1-S10) obtained at the conclusion of the evolution experiment (Week 8) with high (>30-fold) coverage of the very large genome size of *S. arctica*, at 143Mbp. Following very careful variant filtering (Figure 4- Source data 1 and 2), we identified a total of only 26 independently evolved variants with an average of 2.6 mutations per clone (range 1 to 5 per clone) (Figure 4-Source data 3). Of the 26 variants, 24 (∼92.3%) were SNPs (11 coding, and 13 intergenic or intronic), and two were insertions (one coding, one intergenic) (Figure 4-Source data 3). Beside the fact that two of these variants are identical SNPs at the same position (Sarc4_g3900) and have independently evolved in two different backgrounds (Figure 4-Source data 3), no other form of genetic parallelism was detected. While it is very likely that some of these mutations hitch-hiked in the background of beneficial driver mutations, we found that the coding SNP variants were clearly skewed toward non-synonymous changes (9:2), with a cumulative dN:dS-ratio of 1.32. This indicates the general presence of positive selection, and hence adaptative evolution driving this molecular pattern. Based on COG assignments (Figure 4- Source Data 5), four of the changes are orthologous to genes implicated in signal transduction, and two are related to genes with functions in DNA-binding or chromosome condensation.

In the absence of molecular genetic tools, and functional knowledge of many of the mutated gene targets, we set out to better understand how the distinct genetic variants could have influenced sedimentation rates and clump formation by examining the predicted expression dynamics of mutated genes across the cell cycle. The data was derived from a recently published time-resolved transcriptomics dataset of the *S. arctica* life cycle (Dudin et al., 2019). Of the mutated genes, nine showed no expression during the native life cycle, whereas 12 displayed dynamical expression during cellularization, and the remaining five genes were more or less stably expressed (Figure 4 I-K, Figure 4- figure supplement 1K, Figure 4- source data 4). We also annotated all mutation-associated genes based on a recent comprehensive orthology search (Grau-Bové et al., 2017) (Figure 4- source data 5).

The fastest-settling and clumpiest mutant isolated from the population S1, bore a synonymous mutation of a homolog of **i**ncreased **s**odium **t**olerance **1** superfamily (Ist1), and as such likely impacts gene expression rather than gene function. This gene shows a dynamic expression during cellularization and codes for a conserved protein involved in multivesicular body (MVB) protein sorting (Figure 4 I, Figure 4- source data 5) (Dimaano et al., 2008; Frankel et al., 2017). In humans, hIST1 also known as KIAA0174, is a regulator of the endosomal sorting complex required for transport (ESCRT) pathway, and has been shown to be essential for cytokinesis in mammalian cells (Agromayor et al., 2009). Similarly, Ist1 orthologs in both budding and fission yeasts play a role in MVB sorting pathway and, when deleted, exhibit a multiseptated phenotype consistent with a role in cytokinesis and cell separation (Dudin et al., 2017; Xiao et al., 2009).

In the clone derived from population S4, we observed five distinct mutations, two nonsynonymous, one intergenic and two intronic SNPs. Among the two nonsynonymous SNPs, one causes a E90G change in a homolog of human Kanadaptin (SLC4A1AP), which may play a role in signal transduction (Hübner et al., 2002, 2003). Among the non-coding SNPs, one mutation is found in an intron of the 7-dehydrocholesterol reductase (DHCR7), expressed during cellularization and known to be key in the cholesterol biosynthesis pathway, (Fitzky et al., 1998; Prabhu et al., 2016), and the second intronic mutation codes for a STE20-like kinase (SLK) which plays numerous roles in cell-cycle signalling and actin cytoskeleton regulation (Figure 4 J) (Al-Zahrani et al., 2013; Cvrčková et al., 1995; Rohlfs et al., 2007; Y. Wang et al., 2020).

Finally, among the five mutations discovered in the clone derived from population S9, two mutations are in transcription factors that are continually expressed during the cell cycle: an intronic SNP in a basic helix-loop-helix (bHLH) transcription factor, and the sole nonsynonymous SNP leading to A923V change in a gene predicted to encode a nucleotide binding C2H2 Zn finger domain (Fedotova et al., 2017). A third mutation was found in an intron of the highly and dynamically expressed homolog of the **r**egulator of **c**hromosome **c**ondensation **1** (RCC1; Figure 4 K) (Dasso, 1993a; Hadjebi et al., 2008; Qiao et al., 2018). RCC1, is a chromatin-associated protein implicated in several processes including nuclear formation, mRNA splicing and DNA replication (Dasso, 1993b; Forrester et al., 1992; Kadowaki et al., 1992; A. M et al., 1990; O. M et al., 1989). Thus, it may contribute to the accelerated nuclear duplication cycle observed in S9, by impacting cell-cycle progression. Altogether, the mutations identified in both S4 and S9 may affect both cellularization and cell separation.

Among the variants detected in evolved clones with intermediate-settling phenotype, we highlight an intergenic mutation 128bp downstream of Dynamin-1 known to be essential for cytokinesis across different taxa (Konopka et al., 2006; Masud Rana et al., 2013; Rikhy et al., 2015), and a nonsynonymous mutation in a protein similar to Fibrillin-2 (Sarc4_g7365T) which is an extracellular matrix (ECM) glycoprotein essential for the formation of elastic fibres in animals (Figure 4- figure supplement 1K) (M. C. Wang et al., 2009; Yin et al., 2019; Zhang et al., 1994). Altogether, our results across all isolates suggest that a large mutational target affects cellular sedimentation and multicellularity.

### Sedimentation rate variation across *Sphaeroforma* species is driven by cell size and density

Lastly, we examined whether the variation in cell sedimentation observed across different *Sphaeroforma* species (Figure 1E) could also be explained by clumping or increased nuclear number-to-volume ratio. To do so, we investigated the life cycle dynamics, coenocyte volume and nuclear duplication time among distinct *Sphaeroforma* sister species. To date, six different *Sphaeroforma* species have been isolated either in a free-living form or derived from different marine hosts: *S. arctica*, *S. sirkka*, *S. napiecek*, *S. tapetis, S. gastrica* and *S. nootakensis* (Figure 5A) (Ducluzeau et al., 2018; Dudin et al., 2019; Hassett et al., 2015; Jøstensen et al., 2002; W. L. Marshall et al., 2008; W. L. Marshall & Berbee, 2013). Using previously established growth methods for *S. arctica* combined with live and fixed imaging, we first observed that all sister species but *S. tapetis* show a synchronized coenocytic life cycle (Figure 5B, figure 5-figure supplement 1A, Video 4) (Similar observations were also made by A. Ondracka; personal communication). Similar to above mentioned results with fast-settling mutants, sedimentation rate variations (Figure 1E), could be explained by variations in both cell size and cellular density. Indeed, we first observed that both *S. gastrica* and *S. nootakensis* occasionally form clumps, exhibit a lower frequency of cell detachment and thus have an increased cellular density after cell release compared to the other *Sphaeroforma* species (Figures 1C, 5B and C, Video 4). Additionally, despite their increased sedimentation rate (Figure 1C), we found that all sister species exhibited ∼20-45% smaller coenocyte size prior to cell-release when compared to *S. arctica* which reveals an increased cellular density (Figure 5D and figure 5-figure supplement 1B to D, Video 4). Similar to the fast-settling mutants S4 and S9 above, the increase in cellular density was associated with an acceleration of the nuclear division cycles and the subsequent rise in the nuclear number-to-volume ratio (Figure 5D-F and figure 5-figure supplement 1D-F). Notably, *S. sirkka* and *S. napiecek*, both previously isolated as free-living (Ducluzeau et al., 2018; W. L. Marshall & Berbee, 2013), exhibit an increase in nuclear number-to-volume ratio but no ability to form clumps. Altogether, our results show that, similarly to experimentally evolved strains, fast sedimentation variation could occur by both clump formation and/or increase in the nuclear number-to-volume ratio for *Sphaeroforma* species (Figure 5H), and thus might represent in itself a widespread and highly variable phenotype in the marine habitat.

**Figure 5.**
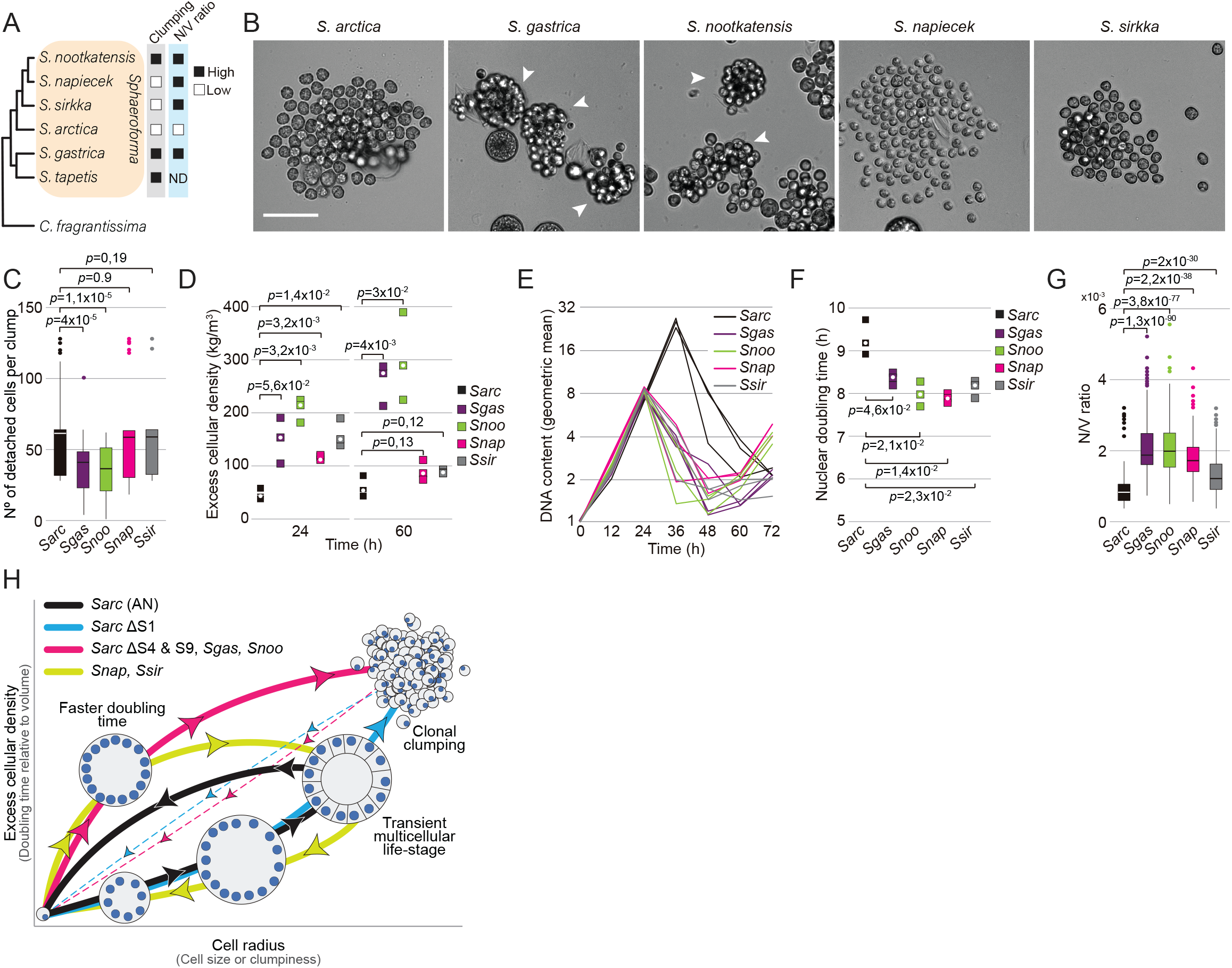
Sedimentation rate variation across *Sphaeroforma* species is associated with clumping and increased nuclear number-to-volume ratio. (A) A cladogram (Hassett et al., 2015), representing the position of all *Sphaeroforma* sister species used in the study and their observed phenotypes relative to *S. arctica*. Cells exhibit either variation in clumpiness and/or nuclear-number to volume ratio (N/C) which is regulated by the doubling time relative to volume. (B) Representative images of different *Sphaeroforma sp.* at cell release. Arrowheads indicate the formation of clumps. Associated with Video 4. Bar, 50µm. (C) Number of cells detaching at cell release, measured from time-lapse movies, show significant differences among the different sister species. (n > 48 coenocytes per *Sphaeroforma sp.*). (D) Excess cellular density of *Sphaeroforma sp.* cells (before cellularization) and clumps (after cell release) at 36 and 72 hours, respectively. Every square represents an independent experiment, and the white circle represents the median. (E) Quantification of mean DNA content per time point for *Sphaeroforma sp.* grown in marine broth at 17°C. Every trace represents an independent experiment (n > 300 coenocytes/timepoint for each independent experiment). (F) Nuclear doubling time, calculated by linear regression of mean nuclear content at time points from 0 hr to 24 hours at 17°C. Every square represents an independent experiment, and the white circle represents the median. (n > 300 coenocytes per timepoint for each independent experiment) (G) Boxplots of nuclear number-to-volume ratio of DAPI stained cells at 17°C for *Sphaeroforma* sister species. Every square represents an independent experiment, and the white circle represents the median. (n > 300 coenocytes per strain). (H) Scheme illustrating a landscape of how phenotypic and genotypic plasticity varies over the physical dimensions of size and cellular density. Grey axis shows the cellular traits affecting these physical dimensions. Colours represent how the different *Sphaeroforma* species investigated in this study behave across their life-cycle. Black = *S. arctica* (AN), Blue = S. *arctica* (S1 mutant), Pink = S. *arctica* (S4 and S9 mutants, *S. gastrica, S. nootakensis*), Green = *S. sirkka, S. napiecek*. Thick lines represent the typical cellular behaviour. Thin and dotted lines indicate that clumping cells may restart the process at any position.

## Discussion

Here, we performed, for the first time, EE on one of the closest unicellular relatives of animals, and we demonstrate that, under suitable selection pressure, the ichthyosporean *Sphaeroforma arctica*, can evolve stable multicellularity. In particular, we observed the independent rise of clump-formation, and faster settling phenotypes across populations within less than 400 generations. The precise detectable onset of the phenotypes varied across lineages and occurred as early as ∼91 generations in lineage S1. We further show that faster sedimentation phenotypes are highly heritable and, according to direct competition assays with the common ancestor, are expected to be adaptive under the environmental conditions of the experiment. Our results add to previous observations of the rapid emergence of multicellularity in yeast and green algae, which all can evolve multicellular clump-forming structures within short evolutionary timescales (Herron et al., 2019; Ratcliff et al., 2012, 2013). As ichthyosporeans proliferate through an uncommon coenocytic life cycle (A. de Mendoza et al., 2015; Dudin et al., 2019; Glockling et al., 2013; W. L. Marshall et al., 2008; W. L. Marshall & Berbee, 2013; Suga & Ruiz-Trillo, 2013), our results show that this evolutionary process of selection for faster sedimentation is accessible at microevolutionary timescales across taxa and organisms with highly diverged modes of proliferation.

In this study, we show that all fast-settling *S. arctica* cells increased their cell size by increasing cell adhesion post-cellularization, leading to the formation of clumps (Figure 5H). Such results are analogous to cell cluster formation in snowflake yeast and *Chlamydomonas reinhardtii* which arises through incomplete separation of mother and daughter cells (Koschwanez et al., 2013; Ratcliff et al., 2012, 2013). Altogether, these results suggest that regulation of sedimentation rate can constrain unicellular species to generate multicellular cell phenotypes by increasing their cell adhesion efficiency. However, we found that ∼31% of the variance could not be explained by cell (clump) size only. Indeed, two fast-settling mutants (S4, S9), exhibited an increase in sedimentation rate prior to clump formation, which was associated with an accelerated nuclear division cell cycle leading to an increase in the number of nuclei per unit volume (Figure 4G and 5H). Previous results have shown that, in *S. arctica*, both nuclear duplication cycles and cell size are uncoupled (Ondracka et al., 2018). Our results support these findings and indicate that nuclear division cycles and cell size could be regulated separately, allowing adaptive change in either, and independently of one another.

By analysing the genomes of evolved isolates, we identified an over-abundance of non-synonymous mutations, indicating positive selection, with at least one and up to five independent mutations in each lineage. Many of the better-characterized genes that carry mutations in either coding or intergenic regions are dynamically expressed during cellularization. Several mutations were found in genes coding for cytoskeletal regulators and cytokinesis proteins which is consistent with previous studies in which cytokinesis deficient mutants were often associated with an incomplete cell separation across taxa (Balasubramanian et al., 1998; Gillmor et al., 2016; Hirono & Yoda, 1997; Huang et al., 2008; Nanninga, 2001). Other mutations were found in genes involved in cell signalling, plasma membrane remodelling and chromatin condensation regulators reflecting a large and accessible mutational target affecting sedimentation rate phenotypes. Cytokinesis defects may lead to incomplete cell separation, which may explain part of the cell clumping phenotype observed in the fast-sedimentors. Given that similar phenotypes emerged independently multiple times with no genetic overlap, we conclude that the mutational target for these traits could be quite large for *S. arctica*, opening the possibility for variation and evolution in multicellularity-related phenotypes.

We found that closely related species exhibit widespread variation in both clump formation and nuclei-to-volume ratio. Given the evolvability of the trait and the few natural samples we examined, it is difficult to argue which cellular states affecting sedimentation rates are ancestral. For example, considering the species tree and the trait of clumpiness (Figure 5A, 5B and Video 4), only three *Sphaerforma* species exhibited this behaviour under our experimental conditions: *S. gastrica*, *S. tapetis* and *S. nootkatensis*. Despite characterizing for the first time all the available isolates of *Sphaeroforma sp.,* the number of studied isolates remains limited (6 species) and thus we cannot distinguish whether this trait is ‘ancestral’ or ‘novel’. Rather, our experimental data suggests that clumpiness and density could evolve over short microevolutionary timescales such as those we have measured in the lab and this in both directions. Evolved lines with highly clumping phenotypes had the competitive edge over their non-clumping common ancestor in relative fitness assays. In contrast, during exponential growth and in the absence of selection, evolved clumpy line were selectively disfavoured relative to their common ancestor, which suggests that clumpy phenotypes are prone to rapid replacement by less clumpy or dense morphs when environments change.

Indeed, marine organisms have evolved various passive or active means of maintaining their position in the water column, for example using motility and/or ingenious approaches to regulate buoyancy (Chen et al., 2019; N. B. Marshall, 1954; Pfeifer, 2015; Strand et al., 2005; Sundby & Kristiansen, 2015; Villareal & Carpenter, 2003). *Sphaeroforma* species are remarkably spherical, immobile, lack flagella and yet exhibit a substantial increase in cell size and density over the life cycle, thus representing a challenge to maintaining buoyancy in marine habitat. This work establishes that *Sphaeroforma*’s cell size and density are subject to tight cellular control and are highly evolvable traits. Taken together, these observations suggest that sedimentation rate is a highly evolvable trait which itself likely shapes the gain and loss of multicellularity.

## Author contributions

O.D. designed the study, performed all the experiments and analysed the data. S.W. analysed and annotated all the genomes from this study. A.M.N. analysed data and models of sedimentation rate. O.D., S.W. and A.M.N. wrote the original draft. O.D. and I.R.T obtained funding. I.R.T supervised the project. All authors reviewed and edited the manuscript.

## Acknowledgments

We thank Macarena Toll Riera, Pierre Gönczy, Gautam Dey, Andrej Ondracka for discussion and comments on the manuscript, Hiroshi Suga, Xavi Grau-Bové for advice on genome analysis, Jon Bråte for cultures of the different *Sphaeroforma* sister species, and Meritxell Antó for technical support. We also thank Andrej Ondracka for sharing his unpublished results on *Sphaeroforma* sister species.

We also acknowledge the CRG Genomics Unit for mRNA library preparation and Illumina sequencing. This work was funded by European Research Council Consolidator Grant (ERC-2012-Co -616960) to I.R.-T.; O.D. was supported by a Swiss National Science Foundation Early PostDoc Mobility fellowship (P2LAP3_171815) a Marie-Sklodowska-Curie individual fellowship (MSCA-IF 746044) and by an Ambizione fellowship from the Swiss National Science Foundation (PZ00P3_185859).

## Methods

### Culture conditions

All *Sphaeroforma sp.* cultures were grown and synchronized as described previously for *Sphaeroforma arctica* (Dudin et al., 2019; Ondracka et al., 2018). Briefly, saturated cultures in Marine Broth (MB) (Difco BD, NJ, USA; 37.4g/L) were diluted into fresh medium at low density (1:250 dilution of the saturated culture) and grown in rectangular canted neck cell culture flask with vented cap (Falcon®; ref: 353108) at either 17°C or 12°C, resulting in a synchronously growing culture. Saturated culture of *Sphaeroforma sp.* are obtained after 3 weeks of growth in MB.

### Experimental evolution

Ten replicate population (S1 to S10) of genetically identical *Sphaeroforma arctica* (AN) were first diluted 250-fold in 5 ml of MB and grown in rectangular canted neck cell culture flask with vented cap (Falcon®; ref: 353108) at 17°C. Cells were grown at 17°C rather than the previously used 12°C in order to increase growth rates and accelerate evolutionary outcome. Every 24 hours, the entire population was transferred into a 15ml falcon tube and allowed to sediment for 2 min on the bench. A 5 ml pipette was then positioned vertically and used to collect 500µl of cell culture from the bottom of the falcon tube. The cells, still vertically positioned in the pipette, were then allowed to sediment once more for 15 second before the transfer of a single drop, equivalent to 20µl, into 5ml of fresh MB (∼250x dilution). Every 7 transfers, a frozen fossil was conserved by adding 10% of DMSO to 1ml of culture and preserved at −80°C. Single clones of each replicate population (S1 to S10) were obtained at the end of week 8 by serial dilutions. Since *S. arctica* grows as a coenocyte, a temporal generation is not defined by a complete coenocytic cycle, but is equivalent to the doubling of the number of nuclei. We estimated the nuclear doubling time by measuring the number of cells and the number of nuclei per cell for each transfer separately (Figure 2- Source data 1). Briefly, the entire EE experiment comprised ∼364 generations, in which all populations underwent a total of 28 complete coenocytic cycles. Each coenocytic cycle included two 24 hours sub-passages, and comprised a total of ∼13 doublings (Figure 2—figure supplement 1A, Figure 2- Source data 1). Importantly, the effective population size was kept in a very narrow range across sub-passages (at ∼10^5^) and thus over the entire experiment (Figure 2—figure supplement 1A, Figure 2- Source data 1). Therefore, in evolutionary terms, the population size was consistently high enough to favor of natural selection over random evolution throughout the course of the experiment. This assumption was reconfirmed genetically by deriving a dN:dS-ratio >1 from sequencing data (see main text).

### Sedimentation rate measurements

Sedimentation rate was measured for *Sphaeroforma sp.* every 12 hours for a total of 72 hours unless indicated otherwise. To ensure reproducibility and homogeneous results, saturated *Sphaeroforma sp.* cultures were sonicated prior to the dilution in fresh MB media (250-fold dilution) using a Branson 450 Digital Sonifier (3 pulses of 15 sec, 10% amplitude). For each measurement, either obtained from different stages of the cell-cycle or from different *Sphaeroforma* species, 1ml of cell culture was added into a disposable plastic spectrophotometer cuvette (semi-micro, 1.5ml) and homogenized by vortex. Optical density (OD_600_) was measured using an Eppendorf® Biophotometer (Model #1631) at T=0, corresponding to the first time point after placing the cuvette in the spectrophotometer. The OD_600_ was then continuously measured every 30 seconds for 3 minutes while cells were slowly sedimenting in the cuvette. To ensure that OD_600_ measurements stayed within the detection limits of the spectrophotometer, early life-stages (T0-T48) were not diluted in the cuvette, whereas later life-stages (T60-T72) were diluted 1/100 in fresh MB media.

For assessing clump dissociation in Figure 2 - figure supplement 1B, AN or S1 cultures were incubated for 2 hours at 37°C in MB, artificial sea water (ASW) (Instant Ocean, 36 g/L) with different salt concentrations ( 18 g/L for 0.5X and 72g/L for 2X) to assess any effect of electrostatic forces, Phosphophate buffered saline (PBS) 1X (Sigma-Aldrich) with either Proteinase K at 200 μg/mL final concentration (New England Biolabs, Ipswich, MA, USA) to assess for any protein-dependent effect or sonication using a Branson 450 Digital Sonifier (3 pulses of 15 sec, 10% amplitude). Only sonication resulted in dissociation of the clumps and a reduction in sedimentation rates.

### Maximum likelihood estimation of sedimentation velocity and cellular density based on OD600 sedimentation rate assay

To briefly summarize, we related our OD600 A.U. sedimentation rate measurements and radius measurements (Figure 1 – Source Data 1) to previously published datasets (SMAYDA & J., 1970) and (Millero & Huang, 2009) (Figure 1 – Source Data 2 and 3) to gain an estimate of our sedimentation rate measurements in metric units, which allowed maximum likelihood estimation of cellular densities based on these estimates (Figure 1 – Source Data 1).

We started by estimating the average radius of cells from perimeter measurements as

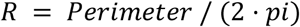

We estimated sedimentation velocity in our measurements by assuming OD600 (OD) changes at a constant proportional rate of change with respect to time t=0 and t=1 by:

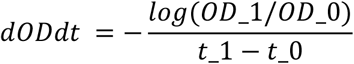

This yielded a rate of change in arbitrary distance units per second which we next sought to relate to metric distance units. For this we turned to the datasets from (Eppley et al., 1967), compiled along with data from other studies by (SMAYDA & J., 1970) in his Appendix Table 1 (our Figure 1 – Source Data 2). In this table, 39 observations included joint values of salt percentages (or seawater density) for measurement, temperature, cell diameters, and sedimentation rate measurements for various phytoplankton isolates.

For each of these observations, we calculated seawater density in experimental assays as *SC + salinity* where salinity constant SC = 35.16504/35 and the salinity is the salt content in *g·l^-1^* (Millero & Huang, 2009). We estimated the specific gravity or density of media used in each sedimentation rate measurement based upon the dataset published by (Millero & Huang, 2009) (Figure 1 – Source Data 3), using a second-order polynomial function of their density measurements as a function of salinity and temperature using R’s *lm()* function. The error of this estimate is extremely low (<4e-3 *kg·m^-3^*) and was not propagated downstream.

We next calculated the density p_p_ of cells across observations based on sedimentation velocity *V* in *m·s*, cell radius *R* in *m* and media density *p_f_* by rearrangement of the terminal velocity equation:

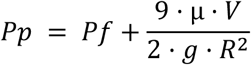

Where dynamic viscosity *µ* = 0.00109 *Pa·s* and gravitational acceleration *g* = 9.780 *m·s^2^*. This yielded a median phytoplankton excess density (p_p_-1000) of 139 *kg·m^-3^* with a range of 30-1300 *kg·m^-3^*. Some excess density estimates exceeding protein (220 *kg·m^-3)^* and cellulose (500 *kg·m^-3^*) have previously been suggested to arise by calcide in diatoms (FOURNIER, 1968). We concluded that this method of estimating cell density yielded similar values to those published by e.g. (FOURNIER, 1968) and (Eppley et al., 1967). For downstream analyses, we excluded obvious outliers, including measurements of samples with densities exceeding that of cellulose, and measurements of *D. rex*, a large diatom which in the Smayda dataset exhibited extraordinarily low sedimentation rates given their size.

To estimate the sedimentation velocities of our dataset, we assumed that our data (Figure 1 – Source Data 1) would fall within the typical measurement in the Smayda dataset (Figure 1 – Source Data 2). For the outlier-excluded subset of the Smayda dataset, we calculated an expected sedimentation velocity for what we would measure in our experimental setup based on the specific gravity of the seawater formulation we used in our measurements (37.4 g/l or a solvent density of 1028.9 *kg·m^-3^*). We then used least-squares minimization to estimate two parameters: a scalar *S* of the AU velocity measurements, *dODdt*, which could best match our dataset with these expected velocities, and an average density parameter *p_p_hat_* which could predict these values and the Smayda dataset’s values based on Stokes’ law. For this we minimized the loss function:

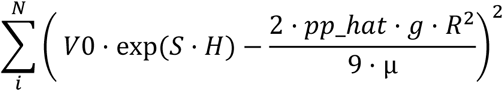

Where *V_0_* is either the seawater-density corrected velocity from Smayda or the *dODdt* parameter we calculated above, and *H* is the one-hot binary scalar in [0,1] corresponding to whether the *i*’th data point in our *N* observations was from Smayda’s or our dataset, respectively. We used R’s *optim()* function with the ‘L-BFGS-B’ method with initial parameters values of *S* = 0.03 and *p_p_hat_* = 100 and repeated this fit for 500 10-fold (10% out-of-bag) bootstrap samples of individual observations across our full dataset and Smayda’s to gain an estimate of the error on the parameters.

*S* and *p_phat_* fits were positively correlated across testing/training folds (Pearson’s ρ = 0.97), however mean values were fairly limited, with *S* = 0.028 ± 0.002 and *p_p_hat_* = 78 ± 6.4 *kg·m^-3^* (with error equal to the standard deviation of estimates across 500 folds). These narrow estimates indicated that the fits were reasonably well-defined by the underlying dataset. The means and standard deviations across predictions of out-of-bag samples are what we have reported as means and standard error in Table 1 and propagated along with inter-replicate and batch error reported in the figures and text.

### Percentage of clumps measurements

To measure percentage of clumps across all mutants and transfers in Figure 2 - figure supplement 1C, we first sonicated saturated *S. arctica* cultures using a Branson 450 Digital Sonifier (3 pulses of 15 sec, 10% amplitude). Cells were then diluted in fresh MB (1:250 dilution) and cell concentration was then measured using a hemocytometer. Approximately 20 cells were then transferred per well in a 96 well plate. Cells were then monitored every 24 hours using an inverted optical microscope and the percentage of clumps observed after cellularization was measured manually using a tally counter. This experiment was performed three independent times and error bars are standard deviations.

### Head-to-head competition

To perform the fitness experiment (head-to-head competition), we used saturated and sonicated cultures of the ancestor (AN), S01 and S03 evolved isolates. Following the measurement of the cell concentration using a neubauer chamber we independently mixed both S01 and S03 with the AN at a ∼1:1 ratio, and subjected them to either the same selective regime as the evolution experiment (see above), or a control culture which was simply maintained in exponential growth without sedimentation selection. The experiment was maintained for 72hrs with passages every 24hrs. To quantify the outcome of the experiment, every 24 hours for 72 hours (7.5 - 12.5 generations), after dilution of the competition culture, we counted the number of clumpy and non-clumpy cells at each passage. For this, a fraction of the cultures was sonicated and highly diluted to obtain fewer than 10 cells per-well of a 96-well plate. These cells were allowed to undergo a full life-cycle in order to count by microscopy the proportion of cells which bore the clumpy phenotype. The high penetrance of the clumpy phenotype in these clones therefore allowed not only a quantification of the clumpy phenotype itself in competition, but also the measurement of overall clonal fitness of these derived isolates.

Each data point for the fitness assay was a count of cells with an associated binomial error. In Figure 2E, Figure 2- figure supplement 1D this error is represented as 95% confidence intervals calculated using the binom.logit() function from the R package “binom” (Sundar Dorai-Raj (2014). binom: Binomial Confidence Intervals for Several Parameterizations. R package version 1.1-1. https://CRAN.R-project.org/package=binom).

The fitness measurements began from highly synchronized cultures, and therefore reporting a per-generation fitness value was clouded by a sudden jump in cellular counts that occurred at 48 hours. Therefore, for simplicity and given the clear time-dependent changes in clumpy phenotype frequencies, we report fitness in terms of log2 fractional change per unit time (“log2 selection coefficient” in days). To obtain these numbers, we first used R’s rbinom() function to resample the dataset 500 times, with each resampling using each sample’s original measurement of clumpy cell count, total cell count, and proportion clumpy. For each resampling, after calculating the new value for proportion clumpy, we logit2 transformed the data and rescaled to the initial measurement with the following equation:

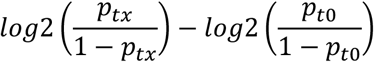

where p is the proportion of the population that was clumpy, the tx underscore is each timepoint, and the t0 underscore is the timepoint of the initial proportion measurement in units days. The slope of the least-squares fit for this quantity with respect to time was obtained by R’s lm() function, yielding the per day change for each resampled biological replicate (the “log2 selection coefficient”). The mean and standard deviation of this coefficient across all resampled replicates were taken as the fitness point estimate and standard error, respectively, and the 95% confidence intervals for Figure 2D is that standard error times 1.96.

### Heritability measurements

To obtain a coarse understanding of how heritable our phenotypes were, we used a statistical genetic approach to quantify the heritability of traits, or what proportion of the variance in phenotype is due to an individual’s inherited state. Briefly, the total Variance of a measured phenotype V_P(Total) can be variance partitioned between Environment, Genetics (or other inherited factors) and technical error or stochastic events (epsilon):

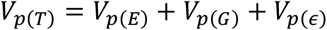

The different partitioning terms in V_P(T) can be expressed as a proportion of V_P(T):

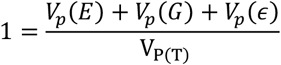

And therefore, we can refer to the proportion of variance due to genetics as pV_P(G).

In the lab, we eliminated environmental variance with tightly controlled experimental conditions such as temperature and the number of hours a measurement was taken after cell cycle synchronization. This allowed us to define heritability therefore as the environment-controlled genotypic variance:

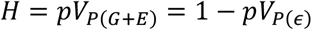

For this, we determined H for DNA content, perimeter length, sedimentation rate and fitness phenotypes by calculating the variance explained by mean phenotype values within distinct genotype/environment combinations (Figure 1 – figure supplement 1A). The results show that H exceeds 95% across phenotypes, and across the entire dataset, H exceeded 99% of the total phenotypic variance (ANOVA F = 1118 on 252 and 735 DF, p = 0). This means that for a typical individual genotype in a given environment, we could predict its average phenotypic measurement with >97% accuracy.

### Microscopy

Microscopy of live and fixed cells was performed using a Zeiss Axio Observer Z.1 Epifluorescence inverted microscope equipped with Colibri LED illumination system and an Axiocam 503 mono camera. An EC Plan-Neofluar 40x/0.75 air objective was used for images of fixed cells and an N-ACHROPLAN 20x/0.45na Ph2 air objective was used for all live imaging, unless indicated otherwise.

### Cell fixation and staining

Throughout this study, saturated *Sphaeroforma* cultures were mildly sonicated prior to diluting them 250X in fresh marine broth to initiate a synchronized culture. To assess for any temperature dependency, cultures were grown at both 17°C and 12°C, and measurements were conducted every 12 hours for a duration of 72 hours. For every time-point, cells were fixed using 4% formaldehyde and 250mM sorbitol for 30 minutes before being washed twice with PBS. For nuclei staining cells were centrifuged at 1000 rpm for 3 min after fixation and washed again three times with PBS before adding DAPI at a final concentration of 5 µg/mL to 5 µl of concentrated sample. DAPI-stained samples were imaged to measure DNA content and coenocyte size. It is important to note that results obtained from fast-settling mutants prior to cell release correspond to measurements of unicellular coenocytes (24 hours for 17°C and 36 hours for 12°C), whereas results collected after cell release correspond to measurements of multi-celled clumps (48 hours for 17°C and 72 hours for 12°C). For cell wall staining, cells were incubated with Calcofluor-white (Sigma-Aldrich) at a final concentration of 5 µg/ml from a 200X stock solution prior to fixation. Cells were then fixed as previously mentioned and concentrated before being disposed between slide and coverslip.

### Live-cell imaging

For live-cell imaging, saturated cultures were diluted 250x in fresh marine broth medium inside a μ-Slide 4 or 8 well slide (Ibidi) at time zero. To ensure oxygenation during the whole period of the experiment, the cover was removed. To maintain the temperature at 17 or 12°C we used a P-Lab Tek (Pecon GmbH) Heating/Cooling system connected to a Lauda Ecoline E100 circulating water bath. To reduce light toxicity, we used a 495nm Long Pass Filter (FGL495M-ThorLabs). For plasma membrane live staining (Figure 3B, Video 3), FM4-64 (Invitrogen) at a final concentration of 10µM from a 100× DMSO diluted stock solution was added at time 0 unless indicated otherwise in figure legends. For cytoplasmic staining of cells in Figure 3- figure supplement 1, cells were either stained with CellTrace™ CFSE Cell Proliferation Kit (Thermofisher) or CellTrace™ Calcein Red-Orange (Thermofisher).

### Image analysis

Image analysis was done using ImageJ software (version 1.52) (Schneider et al., 2012). For nuclear content distribution across *Sphaeroforma sp.’s* life cycle, fixed and DAPI-stained coenocytes were imaged and the number of nuclei per coenocyte was counted using the ObjectJ plugin in imageJ. To compute nuclear duplication times, log2 of geometric mean of DNA content was calculated as: log_2_(geommean) = ∑_*i*_ *f*_*i*_ ∗ log_2_(*x*_*i*_) where fi is the fraction of cells and xi the DNA content (number of nuclei per cell) (ploidy) of each i-th DNA content bin. Nuclear doubling times were computed as linear regression of log2 of geometric mean of DNA content versus time. Note that for *S. tapetis,* nuclear doubling times could not be computed due to the asynchrony in growth. For measurements of cell volume in live and fixed cells we used the oval selection tool to draw the contour of each cell and measured cell perimeter. As cells are spherical, we computed cell volume as: *V* = 4/3*πr^3^* where *r* is the cell radius. For measurements of clumps perimeter, we transformed the images into binaries to ensure later segmentation. We then used the particle analysis function in ImageJ with a circularity parameter set to 0.15–1 to measure cell perimeter. For nuclear number-to-volume ratios, the number of nuclei was divided by the coenocyte volume measured as previously described for fixed cells. All Figures were assembled with Illustrator CC 2020 (Adobe). Several figures were generated using ggplot2 in *R* version 4.0.5 (Wickham, 2016).

### *Sphaeroforma arctica* genome sequencing and assembly

Genomic DNA was extracted for the ancestral strain (AN) and a single clonal isolate from each evolved population (S1-S10) using QIAamp DNA Blood Midi Kit (Qiagen) following the manufacturer’s recommendations from 50 mL culture incubated at 17 °C for 5 days in 75 cm^2^ flasks. The Qubit (*Invitrogen*) quantification ranged between 3 and 13 μg of genomic DNA in total. All of the subsequent steps were performed by the CRG Genomics Unit (Barcelona): sequencing libraries were prepared from the pure high molecular weight DNA using TruSeq DNA HT Library Preparation kit (*Illumina ^®^ HiSeq ^®^ Sequencing v4 Chemistry*). A paired-end library with a target insert size of ∼500 bp was sequenced on an *Illumina^®^* HiSeq2500 platform in paired-end mode, with read lengths of 125 bp. The resulting paired raw read files were demultiplexed by the sequencing facility and data stored in two separate, gzip-compressed FASTQ files of equal sizes. Genome sequencing data has been deposited in NCBI SRA under the BioProject accession PRJNA693121.

### Bioinformatic analyses of the genomes

#### Data processing

On average, each paired-end sequencing library contained ∼58.8 million reads of 125 bp sequence-lengths (Figure 4-Source Data 1), equalling ∼7.35 billion base-pairs (Gbp). From these data, we carefully removed adapter sequences and reads shorter than 50 bp from the raw read data using *trimmomatic* v0.36 (Bolger et al. 2014), yielding an average of ∼4.36 Gbp of filtered sequence data per genome (Figure 4-Source Data 1). The quality of both raw and trimmed sequencing data was assessed in FastQC v0.11.7 (Andrews, 2010). In more detail, based on the FastQC output for raw reads, we initiated trimming by calling the following parameters:

*ILLUMINACLIP:Nextera+TruSeq3-PE2.fa:3:25:10 CROP:110 LEADING:30 TRAILING:25 SLIDINGWINDOW:4:28 MINLEN:50*

This translates into the following trimming steps:

- cut adapters and other *Illumina* specific sequences using a combined file of default adapters (“Nextera.fa” AND “TruSeq3-PE2.fa”) to catch as many spurious contaminations during library prep as possible, with seed mismatches = 3; palindrome clip threshold = 25; simple clip threshold = 10;
- end-clipping of the final 15bp of all reads, due to evidence for elevated adapter content;
- quality-clip all bases on leading ends as long as bases were of lower quality than Q < 30;
- removing all bases on trailing ends as long as bases were of lower quality than Q < 25; and,
- finally, conducting a sliding window approach, where the reads were trimmed once the average quality within a window of four consecutive bases falls below a threshold of Q <28.

The trimming output consisted of four FASTQ files for each genome, of which two files contained intact paired-end reads, and another two files containing all unpaired reads for each end separately.

#### Repeat-masking of reference genome

We relied on the latest (i.e., fourth) assembly version of the *S. arctica* reference genome (Sarc4; (Dudin et al., 2019)) for variant detection and annotation. Initial runs of the read-mapping steps revealed a high proportion of tightly clustered variants (∼20.4%) concentrated in certain regions of the genome (Figure 4-Source Data 2). We investigated this phenomenon and determined that the issue was caused by repetitive stretches of sequence and thus decided to mask all of the potentially problematic repeat regions. For this, repetitive regions were screened for and properly annotated in *RepeatMasker* v4.1.1 (Smit et al., 2015) relying on the 20181026 release of *GIRI RepBase* database for annotations (Bao et al., 2015) and applying the slow high-sensitivity search mode with the following parameters: *-pa 10 -s -gff -excln -species Opisthokonta*.

#### Variant prediction

We performed read alignment, variant calling, variant filtering in CLC Genomic Workbench v20.0.4 (^©^*Qiagen*) and our analytical pipeline was structured into the following steps:

- *<u>Read alignment</u>.* Paired-end reads were merged (assuming insert lengths between 400 and 600bp), and both paired and unpaired reads were aligned against the reference genome (with settings: Length fraction ≥ 0.9; Similarity fraction ≥ 0.9; Match score = 1; Mismatch cost = 3; Insertion/Deletion Open Cost = 5; Insertion/Deletion extend cost = 3; Global Alignment = no). Finally, reads were deduplicated (maximum representation of minority sequence = 0.2).
- *<u>Variant calling</u>.* We called variants using CLC’s “Fixed Ploidy Variant Detection” assuming a haploid genome (Ploidy = 1) for *Sphaeroforma arctica*. We further required a variant probability of at least 90%, ignoring positions in excess of 90x coverage, broken read pairs and non-specific matches. All variants needed to be covered by at least 10 variant-bearing reads, and a minimum consensus of 80% (i.e., at least 10/12 variant supporting reads). We also applied the following read quality filters: neighborhood radius = 10; minimum central quality = 20; minimum neighborhood quality = 20), and read direction filters (direction frequency = 0.05; relative read direction filter = yes (significance = 0.05); read position filter = yes (significance = 0.05).
- Variant filtering. 1) To consider only variants that have emerged through the course of evolution, we automatically removed mutations present in the ancestor (AN). 2) We then went on to manually curate all mutation predictions. We did this by aligning the mapping tracks (profiles) of all re-sequenced genomes (AN, S1 – S10) and screening all initial candidate mutations. The overwhelming majority of variant calls were visually shared across all evolved clones but not called universally due to extremely low statistical support, or low local coverage in some of the genomes. The final dataset hence only contained 25 predicted variants (one shared among two lineages, hence, representing 26 independent mutations) and exported these for each of the ten evolved single clones as Variant Call File (VCF) format.
- *<u>Variant annotation</u>.* We annotated filtered variants from converted VCF files with *breseq v0.33.2* (Deatherage & Barrick, 2014) for convenience. More specifically, we converted VCF files into *breseq’s* Genome Diff (GD) file format using the command “*gdtools VCF2GD*”. We then annotated all genomes in GD format jointly by running the command “*gdtools ANNOTATE*” and specifying original *S. arctica* genome assembly (Sarc4) in GenBankFormat (GBK) as reference. Mutations were tabulated and sorted. Finally, we counted and categorized both observed mutations and non-synonymous and synonymous sites at risk across the reference genome using the command “*gdtools COUNT -b*” for statistics and the calculation of the compound dN/dS-ratio.

---

**Figure 1. figure supplement 1.**
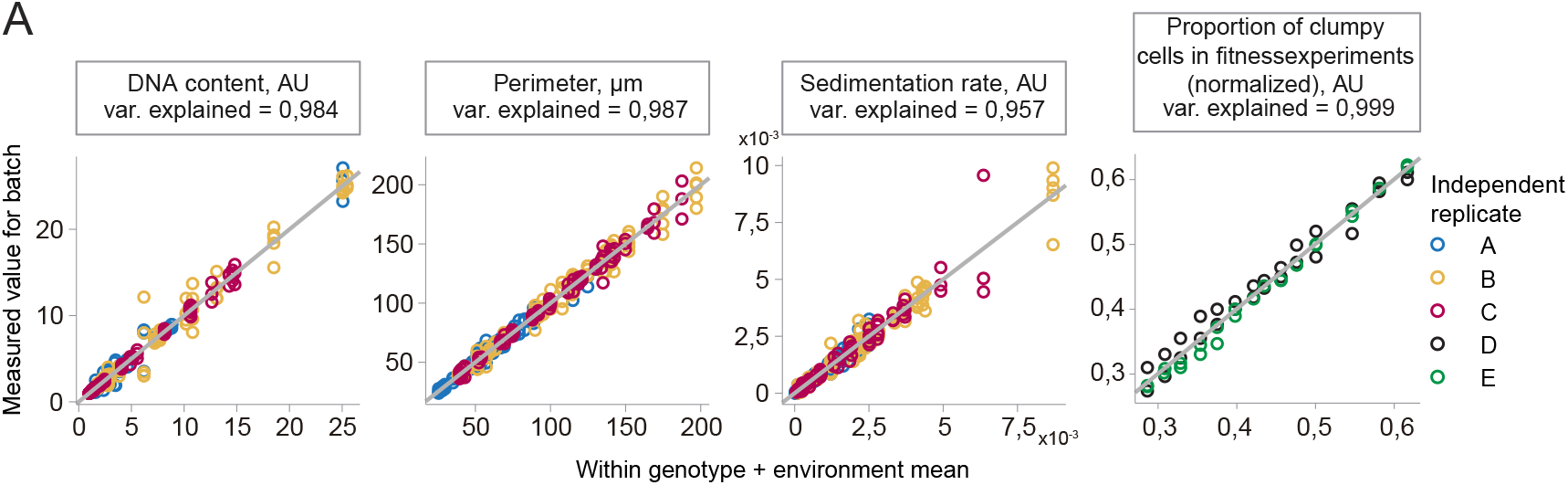
Variance measurements and heritability of the characterised phenotypes across this study. (A) Measurements of technical error or heritability of traits show that the phenotypes measured for the evolved isolates were highly heritable. Arithmetic mean values for four key phenotypes were calculated for each genotype + environment combination (horizontal X axis) and plotted against individual measured values (vertical Y axis) across replicates performed on separate days (“batches” - colours). Environments included different temperatures, and different timings after cell cycle synchronization. Horizontal facet strips indicate the phenotype, its unit of measure, and the heritability of the trait, i.e. the variance explained by the mean genotype + environment value. Grey lines represent where X = Y.

**Figure 1 – Source Data 1. Our data used for estimating sedimentation rate in metric units and density in mass per volume.**

’V_au’ is sedimentation rate in AU OD600 units per second.

’R_meters’ is the mean radius of cells for the genotype and environment

’Species’, genotype, temp, hours_growth are information for aggregating the data based on genotype and environment.

’V_mu’ is the maximum likelihood estimate of cellular sedimentation rate - the mean of all out-of-bag samples used in fitting - in meters per second.

’V_se’ is the standard deviation of cellular sedimentation rate across all out-of-bag samples in the fitting.

’pp_mu’ the maximum likelihood estimate of density - the mean of all out-of-bag samples used in fitting - in kg/m^3^.

’pp_se’ the maximum likelihood estimate of density - the standard deviation of all out-of-bag samples used in fitting - in kg/m^3^.

**Figure 1 – Source Data 2. Smayda dataset used for calibrating our dataset. Adapted from (SMAYDA & J., 1970) Appendix Table 1**

’classification’, species are from his table annotation.

’salt_percent’ is the percent salt reported in the table.

’V_meters_per_second’ is the velocity reported, converted from meters per day to meters per second.

’Pf’ is the density of the media based on the reported temperature and salt concentration, estimated by the data from Millero and Huang.

’R_meters’ is the mean radius in meters. When a range was given, this is the average of that range.

’Pp_’ is the density of the sample in kg/m^3^

’Pp’ is the excess density kg/m^3^

’V_exp’ is the expected velocity in the seawater used in our experiments (37.4 g/l)

**Figure 1 – Source Data 3. The Millero and Huang data (Millero & Huang, 2009) used for estimating seawater density based on salinity and temperature.**

’temp_C’ temperature in degrees Celsius ‘salinity’ salt in g/l

’density_kgm3’ excess density in kg/m^3^

’density’ density in kg/m^3^

**Figure 1 – Source Data 4. Fitness experiment data used for heritability measurements. ‘batch’ is an arbitrary alphabetical letter referring to the date the replicate was performed (either D or E)**

‘rep’ is the replicate within batch/day (rep1 or rep2)

‘geno’ is the genotype information (which genotype was being competed against the “AN” ancestor, or “alone” if alone in competition)

‘time’ is the time the sample was collected in units’ hours

‘sel_cond’ is the condition of the selection - either the initial population (“init”), no selection for sedimentation rate (“NO_SEL”), selection for sedimentation rate (“YES_SEL”)

‘NOT_CLUMPY’ is the count of microcolonies with not clumpy phenotype

‘YES_CLUMPY’ is the count of microcolonies with clumpy phenotype

‘cell_density’ is the measured cell density at time of dilution after period of growth

‘N’ is the total population growth calculated from cell_density per fold dilutions (1/250 per day).

‘gen’ is the number of generations of growth based on cellular counts.

Note for both ‘N’ and ‘gen’ that due to the synchronized coenocytic growth the cultures, they underwent a large increase in cell count after 48 hours.

‘date’ is the date the replicate was performed (April or May 2019)

‘temp’ is the temperature of the competition.

**Figure 2- figure supplement 1.**
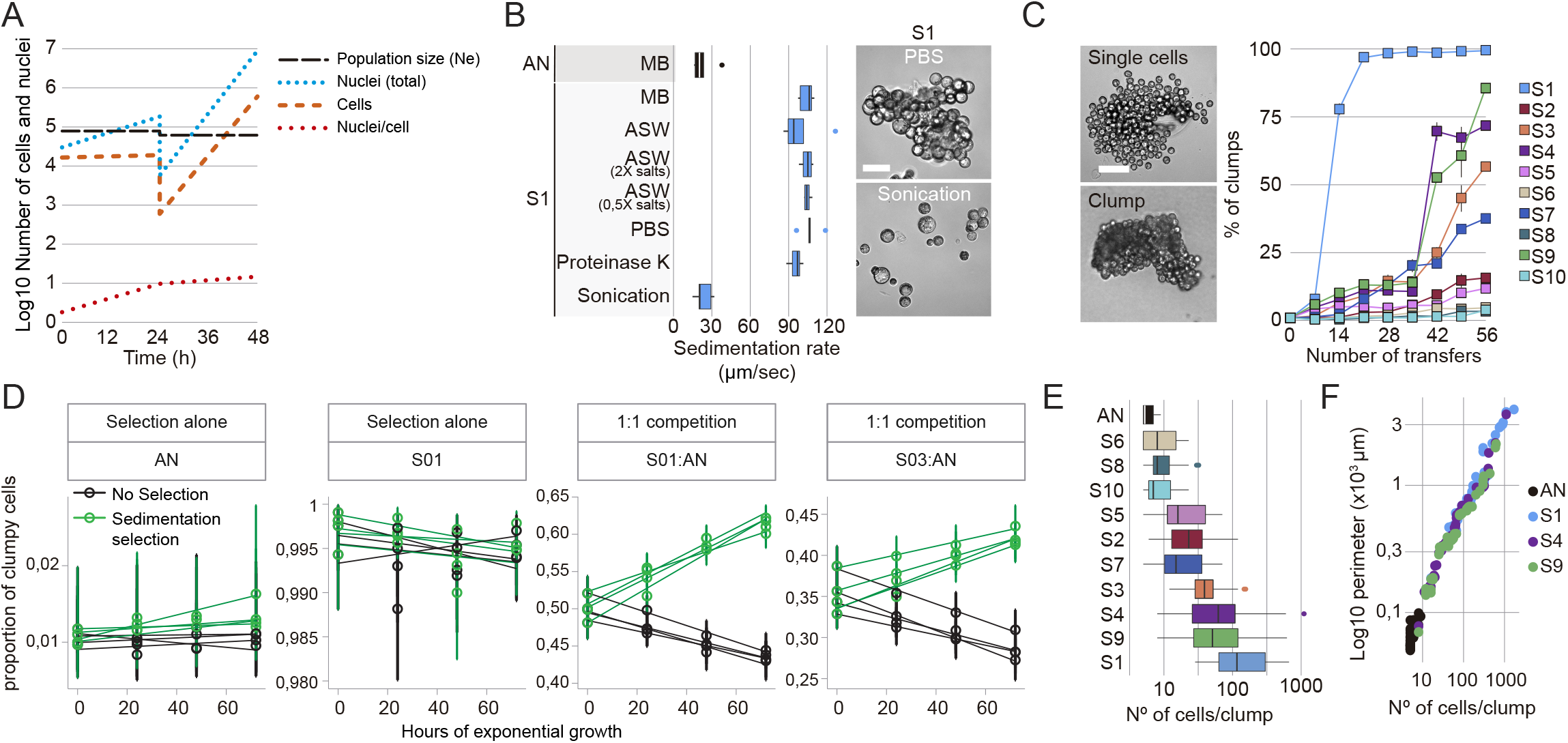
Clump size of evolved mutants correlates with increased number of cells per clump. (A) Approximation of the constant population size derived from nuclear doublings over the course of 48 hours. Nuclei numbers were estimated from total cell number and average number of nuclei per cells (after cell-sorting). At 24 hours, a constant fraction of cells has been transferred to fresh medium, which induces a bottleneck. (B) Sedimentation rates of S1 clumps after incubation for two hours in different media or after sonication. MB= Marine Broth, ASW = Artificial sea water (Salt concentration = 36.4 g/L), 2X (Salts concentration = 72.8g/L), 0.5X (Salts concentration = 18.2g/L), PBS = Phosphate Buffered Saline 1X, Proteinase K 200 μg/mL and sonication (3 pulses of 15 sec, 10% amplitude). Representative images on the right show the dissociation of S1 clumps after sonication. Bar, 50µm. (C) Percentage of clumps formed after 72h of all evolved mutants per number of transfers (n > 200 coenocytes at cell-release per strain). Example images on the left of detached or clumpy cells after cell-release. Bar, 50µm. (D) Raw data for the frequency with respect to time of each independent replicate of strains Ancestor (AN) and clumpy evolved isolates S01 and S03 in head-to-head competitions with the ancestral strain (Two days, two replicates per day). Each point represents a measurement of a given replicate at the given timepoint under the two selection regimes (colours – Green = in presence of sedimentation rate selection, Black = in absence of selection), and error bars are 95% confidence intervals. Lines are least-squares regressions of each replicate across timepoints to guide the eye. (E) Number of cells per clump in all evolved mutants illustrates the linear correlation between sedimentation rate and number of cells per clump. (n = 13 clumps for AN and 50 clumps for all evolved mutants of at least 5 attached cells together). (F) Linear correlation between clump size of the fast-settling mutants (S1, S4, S9) and number of cells per clump compared to the ancestral strain (AN).

**Figure 3- figure supplement 1.**
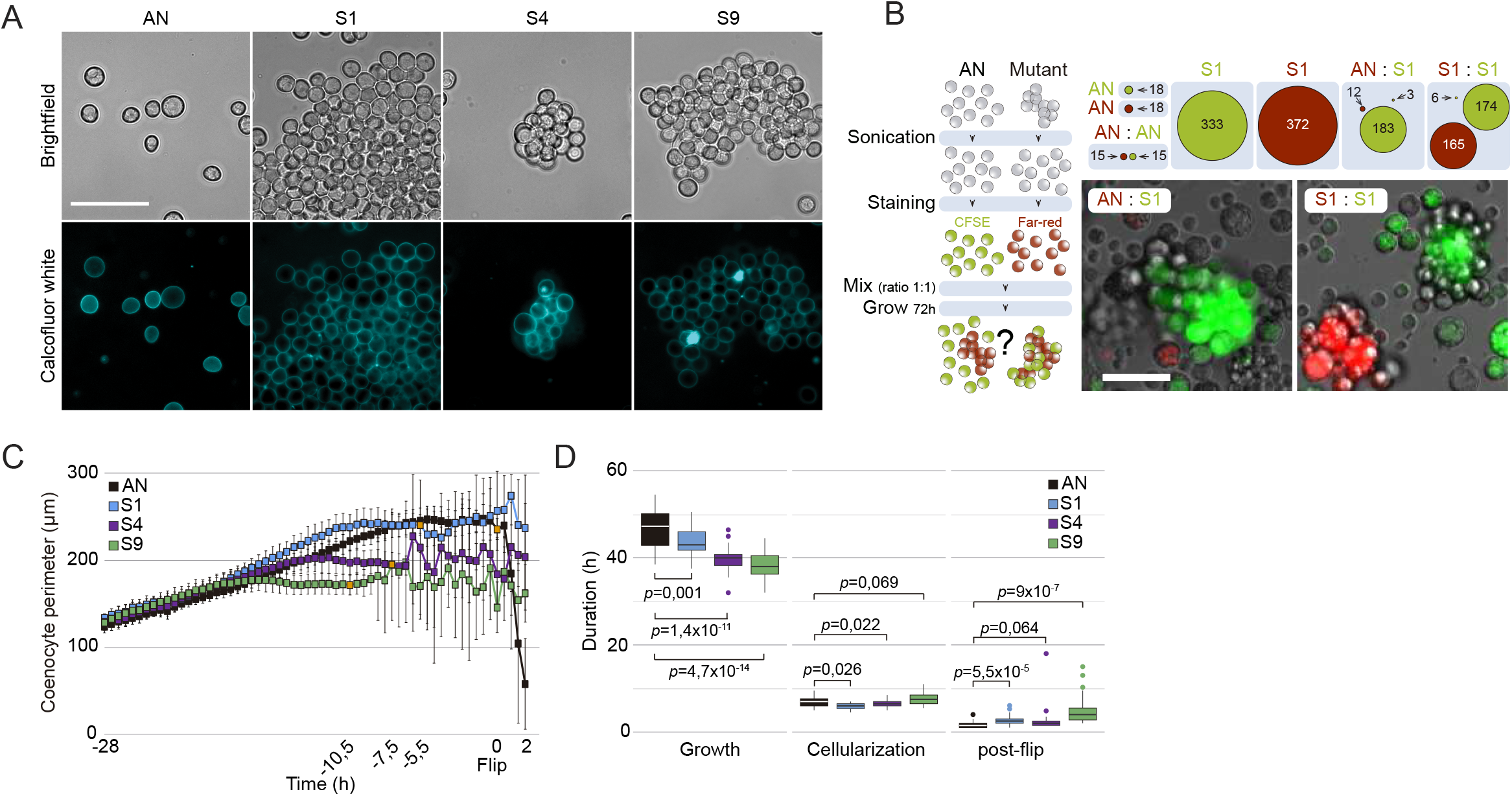
Fast-settling mutants cannot form clumps by aggregation and show discreet differences in life-stages duration. (A) Cell-wall staining show presence of a separating cell wall between individual cells in the clumps of all fast-settling mutants. Bar, 50µm. (B) Experimental design and measurements of S1 clump formation by aggregation. AN and S1 cells are separated by sonication and stained with different cellular dyes before being mixed together for a complete life cycle of 72 hours. Clumps of each separate or mixed colours are then counted by microscopy. Circles represent the number of clumps observed in each condition (more than 5 cells attached together). We show that when S1 is mixed with either AN or itself (1:1 ratio) it mostly forms mono-coloured clumps. Representative images of mono-coloured clumps. Bar, 50 µm (C) Mean coenocyte perimeter over time (10 cell traces per strain) at 12°C, aligned to time 0, reveals discreet differences in coenocyte perimeter and life-cell stages among fast-settling mutants. Orange squares represents the flip timepoint in each trace. (D) Duration of growth, cellularization and post-flip represented as box-plots at 12°C (n > 28 coenocytes each).

**Figure 4- figure supplement 1.**
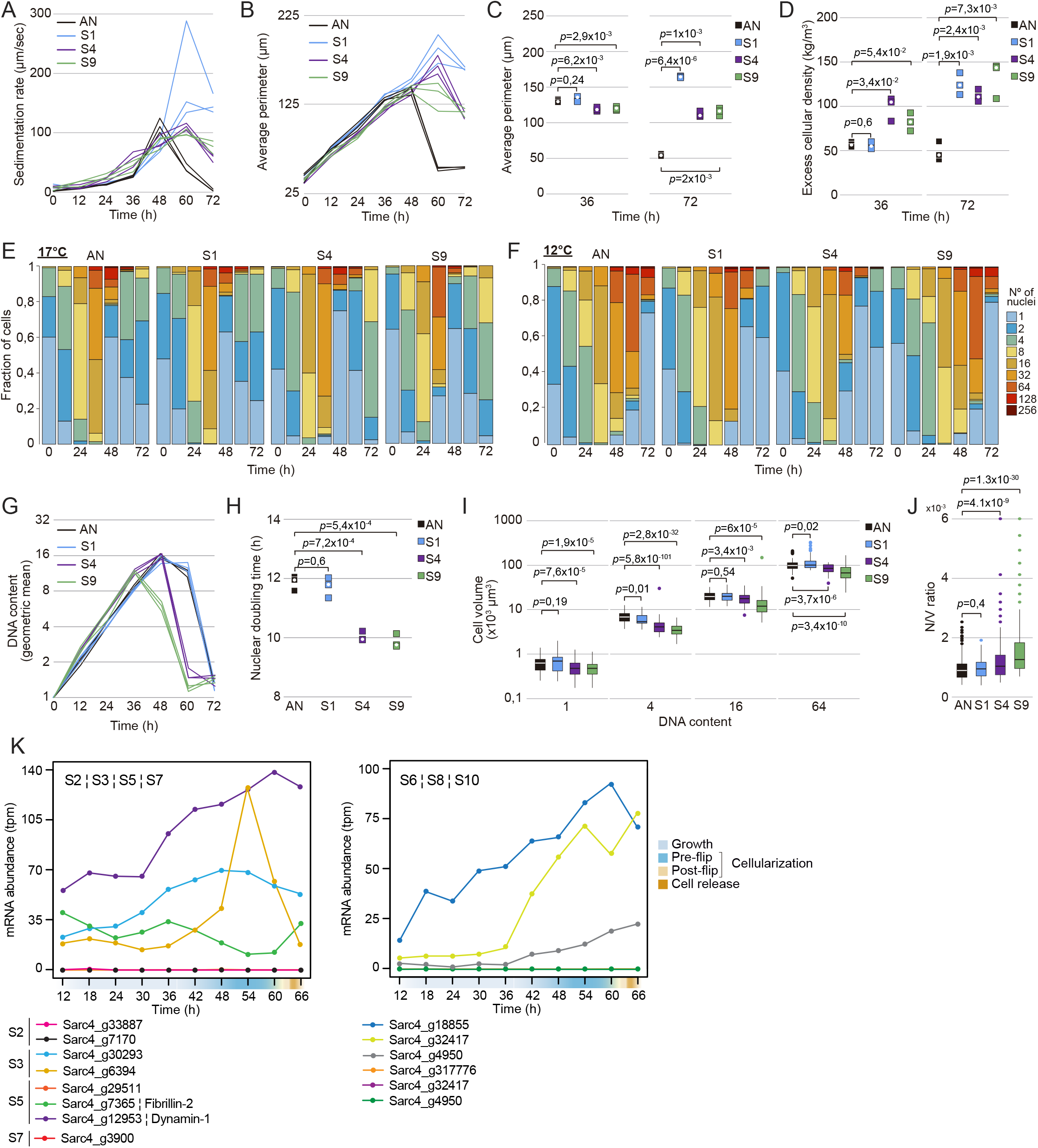
Increased sedimentation rates in fast-settling mutants is independent of temperature. (A) Sedimentation rates of *S. arctica* AN and evolved mutants during the life cycle at 12°C. Every trace represents an independent experiment. (B) Average perimeter measured from fixed cells every 12 hours over a complete life cycle of 72 hours at 12°C shows that fast-settling mutant increase their size upon-cell release. Every trace represents an independent experiment (n > 130 measurements per timepoint for each independent experiment). (C) Average perimeter of fast-settling cells and clumps at 36 and 72 hours respectively show that S4 and S9 single cells and clumps have a smaller size when compared to S1 at 12°C. Every square represents an independent experiment, and the white circle represents the median. (D) Excess cellular density of fast-settling mutants (before cellularization) and clumps (after cellularization) at 36 and 72 hours, respectively. Every square represents an independent experiment, and the white circle represents the median. (E) Distributions of nuclear content of *S. arctica* AN and fast-settling mutants during the life cycle at 17°C measured by microscopy of DAPI-fixed cells (n > 400 coenocytes per timepoint). (F) Distributions of nuclear content of *S. arctica* AN and fast-settling mutants during the life cycle at 12°C measured by microscopy of DAPI-fixed cells (n > 420 coenocytes per timepoint for each independent experiment). (G) Quantification of mean DNA content per time point for fast-settling mutants grown in marine broth at 12°C. Every trace represents an independent experiment (n > 420 coenocytes per timepoint for each independent experiment) (H) Nuclear doubling time, calculated by linear regression of mean nuclear content at time points from 0 to 24 hours at 12°C. Every square represents an independent experiment, and the white circle represents the median (n > 420 coenocytes per timepoint for each independent experiment). (I) Boxplots of cell volume measurements of DAPI-stained fixed cells at 12°C. For 1-, 4-, 16-, and 64-nuclei cells. Cells with one nucleus represent new-born cells at the end of the experiment (n > 50 coenocytes per DNA content). (J) Boxplots of nuclear number-to-volume ratio of DAPI stained cells at 12°C. Every square represents an independent experiment, and the white circle represents the median (n > 300 coenocytes per strain). (K) Temporal transcript abundance of genes mutated in intermediate and slow-settling phenotypes across the native life cycle of *S. arctica*.

**Figure 5- figure supplement 1.**
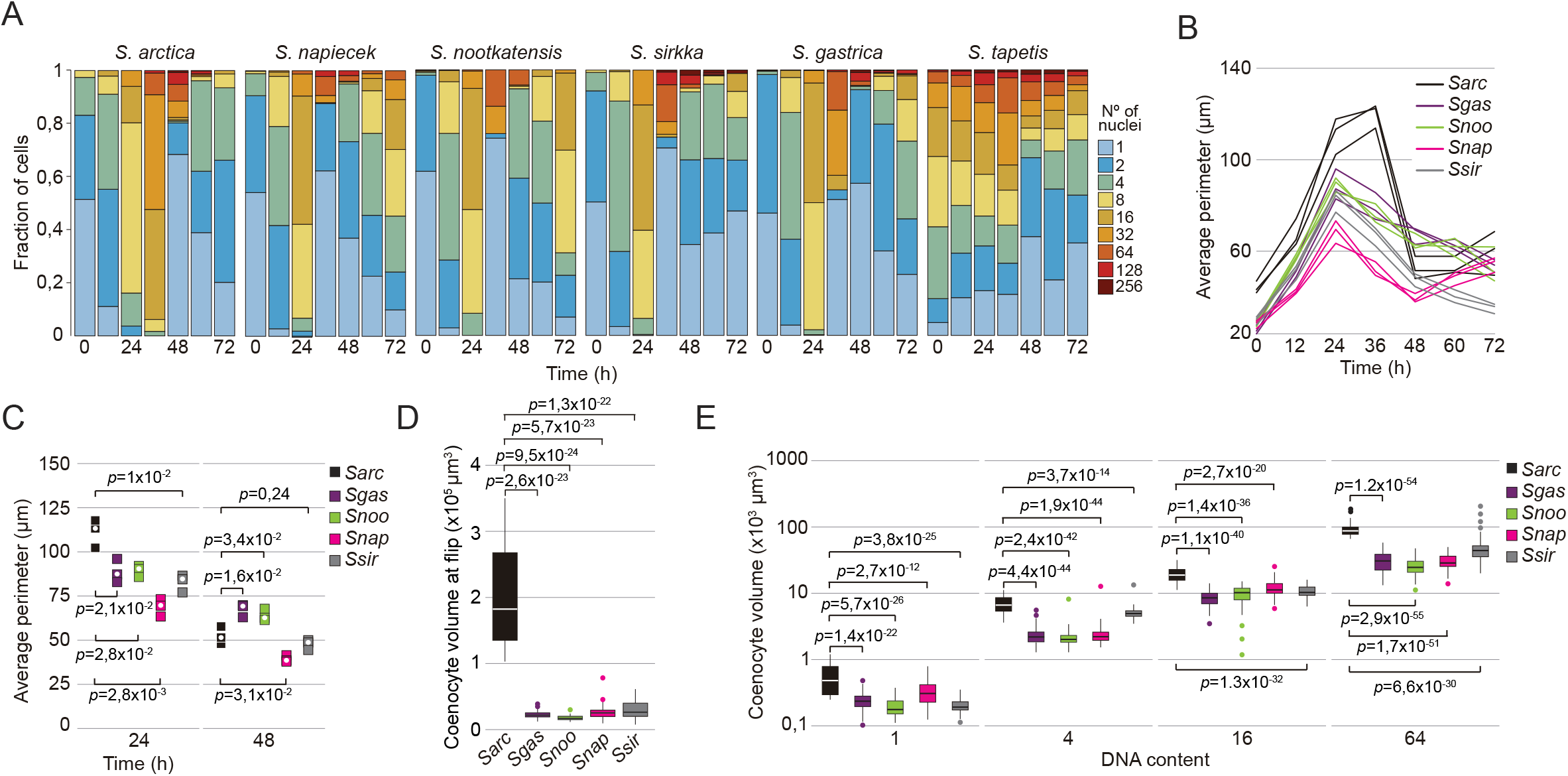
*Sphaeroforma* sister species are distinct in growth, sedimentation rates and cell volume. (A) Distributions of nuclear content of *Sphaeroforma* species during the life cycle at 17°C of DAPI-fixed cells measured by microscopy. Note that *S. tapetis* is asynchronous compared to all other species (n > 300 coenocytes per timepoint for each independent experiment). (B) Average perimeter measured from fixed cells every 12 hours over a complete life cycle of 72 hours at 17°C of *Sphaeroforma sp.* Every trace represents an independent experiment (n > 140 measurements per timepoint for each independent experiment). (C) Average perimeter of *Sphaeroforma sp.* cells and clumps at 0, 24, and 48 hours, respectively, show that *S. gastrica, S. nootakensis, S. napiecek* and *S. sirkka* are smaller in size at 17°C. Every square represents an independent experiment, and the white circle represents the median (n > 140 measurements per timepoint for each independent experiment). (D) Coenocyte volume at flip, measured from time-lapse movies, show that all *Sphaeroforma* species apart from *S. arctica* have significantly smaller coenocyte volume at flip (n > 50 coenocytes per strain). (E) Boxplots of cell volume measurements of DAPI-stained fixed *Sphaeroforma sp.* coenocytes at 17°C. For 1-, 4-, 16-, and 64-nuclei cells. Cells with one nucleus represent new-born cells at the end of the experiment. (n > 50 coenocytes per DNA content).

**Video 1. Video of *Sphaeroforma arctica* AN and fast-settling (S1, S4, S9) cultures sedimenting.**

Time interval between frames is 0.5 sec. The movie is played at 7 frames per second (fps). We can observe rapid cell sedimentation in S1, S4 and S9 when compared to AN. Note that S1 clumps are bigger and sediment faster than the two other mutants. The movie was acquired for cultures pre-grown for 72 hours at 12°C, and obtained with a mobile phone (Samsung A20).

**Video 2. Time-lapse video of synchronized cells of *S. arctica* AN and fast-settling mutants (S1, S4, S9).**

Time interval between frames is 30 min. The movie is played at 7 fps. Four distinct cells can be seen undergoing a full life cycle at 12°C with the release of detached new-born cells for AN or clumps for the mutants. Bar, 50 µm.

**Video 3 Time-lapse video of cells of *S. arctica* AN and fast-settling mutants (S1, S4, S9) stained with the membrane dye (Fm4-64).**

Time interval between frames is 15 min. The movie is played at 7fps. Clumps can be seen being formed after plasma membrane invaginations followed by cell release. Obtained with an epifluorescence microscope. Bar, 50 µm.

**Video 4. Time-lapse video of six different *Sphaeroforma* species undergoing a complete life cycle.**

Time interval between frames is 30 min. The movie is played at 7 fps. Note the asynchrony of *S. tapetis*, the capacity to clump of *S. gastrica* and *S. nootakensis,* and the small cell size of all *Sphaeroforma* species compared to *S. arctica*. Bar, 50 µm.

## References

Agromayor, M., Carlton, J. G., Phelan, J. P., Matthews, D. R., Carlin, L. M., Ameer-Beg, S., Bowers, K., & Martin-Serrano, J. (2009). Essential role of hISTI in cytokinesis. Molecular Biology of the Cell. https://doi.org/10.1091/mbc.E08-05-0474

Al-Zahrani, K. N., Baron, K. D., & Sabourin, L. A. (2013). Ste20-like kinase SLK, at the crossroads: A matter of life and death. In Cell Adhesion and Migration. https://doi.org/10.4161/cam.22495

Alegado, R. A., Brown, L. W., Cao, S., Dermenjian, R. K., Zuzow, R., Fairclough, S. R., Clardy, J., & King, N. (2012). A bacterial sulfonolipid triggers multicellular development in the closest living relatives of animals. ELife. https://doi.org/10.7554/eLife.00013

Allen, W. E. (1932). Problems of Flotation and Deposition of Marine Plankton Diatoms. Transactions of the American Microscopical Society, 51(1), 1. https://doi.org/10.2307/3222044

Andrews, S. (2010). FastQC - A quality control tool for high throughput sequence data. http://www.bioinformatics.babraham.ac.uk/projects/fastqc/. Babraham Bioinformatics.

Balasubramanian, M. K., Mccollum, D., Chang, L., Wong, K. C. Y., Naqvi, N. I., He, X., Sazer, S., & Gould, K. L. (1998). Isolation and Characterization of New Fission Yeast Cytokinesis Mutants.

Bao, W., Kojima, K. K., & Kohany, O. (2015). Repbase Update, a database of repetitive elements in eukaryotic genomes. Mobile DNA. https://doi.org/10.1186/s13100-015-0041-9

Beardall, J., Allen, D., Bragg, J., Finkel, Z. V., Flynn, K. J., Quigg, A., Rees, T. A. V., Richardson, A., & Raven, J. A. (2009). Allometry and stoichiometry of unicellular, colonial and multicellular phytoplankton. New Phytologist, 181(2), 295–309. https://doi.org/10.1111/j.1469-8137.2008.02660.x

Bonner, J. T. (1998). The origins of multicellularity. Integrative Biology: Issues, News, and Reviews, 1(1), 27–36. https://doi.org/10.1002/(SICI)1520-6602(1998)1:1<27::AID-INBI4>3.0.CO;2-6

Brunet, T., & King, N. (2017). The Origin of Animal Multicellularity and Cell Differentiation. In Developmental Cell. https://doi.org/10.1016/j.devcel.2017.09.016

Brunet, T., Larson, B. T., Linden, T. A., Vermeij, M. J. A., McDonald, K., & King, N. (2019). Light-regulated collective contractility in a multicellular choanoflagellate. Science (New York, N.Y.). https://doi.org/10.1126/science.aay2346

Chen, M., Tian, L. L., Ren, C. Y., Xu, C. Y., Wang, Y. Y., & Li, L. (2019). Extracellular polysaccharide synthesis in a bloom-forming strain of Microcystis aeruginosa: implications for colonization and buoyancy. Scientific Reports. https://doi.org/10.1038/s41598-018-37398-6

Cvrčková, F., De Virgilio, C., Manser, E., Pringle, J. R., & Nasmyth, K. (1995). Ste20-like protein kinases are required for normal localization of cell growth and for cytokinesis in budding yeast. Genes and Development. https://doi.org/10.1101/gad.9.15.1817

Dasso, M. (1993a). RCC1 in the cell cycle: the regulator of chromosome condensation takes on new roles. Trends in Biochemical Sciences. https://doi.org/10.1016/0968-0004(93)90161-F

Dasso, M. (1993b). RCC1 in the cell cycle: the regulator of chromosome condensation takes on new roles. Trends in Biochemical Sciences, 18(3), 96–101. https://doi.org/10.1016/0968-0004(93)90161-F

de Mendoza, A., Suga, H., Permanyer, J., Irimia, M., & Ruiz-Trillo, I. (2015). Complex transcriptional regulation and independent evolution of fungal-like traits in a relative of animals. ELife, 4. https://doi.org/10.7554/eLife.08904

Deatherage, D. E., & Barrick, J. E. (2014). Identification of mutations in laboratory-evolved microbes from next-generation sequencing data using breseq. Methods in Molecular Biology. https://doi.org/10.1007/978-1-4939-0554-6_12

Dimaano, C., Jones, C. B., Hanono, A., Curtiss, M., & Babst, M. (2008). Ist1 regulates Vps4 localization and assembly. Molecular Biology of the Cell. https://doi.org/10.1091/mbc.E07-08-0747

Ducluzeau, A.-L., Tyson, J. R., Collins, R. E., Snutch, T. P., & Hassett, B. T. (2018). Genome Sequencing of Sub-Arctic Mesomycetozoean Sphaeroforma sirkka Strain B5, Performed with the Oxford Nanopore minION and Illumina HiSeq Systems. Microbiology Resource Announcements. https://doi.org/10.1128/mra.00848-18

Dudin, O., Merlini, L., Bendezú, F. O., Groux, R., Vincenzetti, V., & Martin, S. G. (2017). A systematic screen for morphological abnormalities during fission yeast sexual reproduction identifies a mechanism of actin aster formation for cell fusion. PLoS Genetics, 13(4). https://doi.org/10.1371/journal.pgen.1006721

Dudin, O., Ondracka, A., Grau-Bové, X., Haraldsen, A. A. B., Toyoda, A., Suga, H., Bråte, J., & Ruiz-Trillo, I. (2019). A unicellular relative of animals generates a layer of polarized cells by actomyosin-dependent cellularization. ELife, 8. https://doi.org/10.7554/eLife.49801

Elena, S. F., & Lenski, R. E. (2003). Evolution experiments with microorganisms: the dynamics and genetic bases of adaptation. Nature Reviews Genetics, 4(6), 457–469. https://doi.org/10.1038/nrg1088

Eppley, R. W., Holmes, R. W., & Strickland, J. D. H. (1967). Sinking rates of marine phytoplankton measured with a fluorometer. Journal of Experimental Marine Biology and Ecology, 1(2), 191–208. https://doi.org/10.1016/0022-0981(67)90014-7

Fairclough, S. R., Dayel, M. J., & King, N. (2010). Multicellular development in a choanoflagellate. In Current Biology. https://doi.org/10.1016/j.cub.2010.09.014

Fedotova, A. A., Bonchuk, A. N., Mogila, V. A., & Georgiev, P. G. (2017). C2H2 zinc finger proteins: The largest but poorly explored family of higher eukaryotic transcription factors. In Acta Naturae. https://doi.org/10.32607/20758251-2017-9-2-47-58

Finkel, Z. V., Beardall, J., Flynn, K. J., Quigg, A., Rees, T. A. V., & Raven, J. A. (2010). Phytoplankton in a changing world: Cell size and elemental stoichiometry. In Journal of Plankton Research (Vol. 32, Issue 1, pp. 119–137). Oxford Academic. https://doi.org/10.1093/plankt/fbp098

Fisher, R. M., Cornwallis, C. K., & West, S. A. (2013). Group formation, relatedness, and the evolution of multicellularity. Current Biology, 23(12), 1120–1125. https://doi.org/10.1016/j.cub.2013.05.004

Fitzky, B. U., Witsch-Baumgartner, M., Erdel, M., Lee, J. N., Paik, Y. K., Glossmann, H., Utermann, G., & Moebius, F. F. (1998). Mutations in the Δ7-sterol reductase gene in patients with the Smith-Lemli-Opitz syndrome. Proceedings of the National Academy of Sciences of the United States of America. https://doi.org/10.1073/pnas.95.14.8181

Forrester, W., Stutz, F., Rosbash, M., & Wickens, M. (1992). Defects in mRNA 3’-end formation, transcription initiation, and mRNA transport associated with the yeast mutation prp20: possible coupling of mRNA processing and chromatin structure. Genes & Development, 6(10), 1914–1926. https://doi.org/10.1101/GAD.6.10.1914

Fournier, R. O. (1968). OBSERVATIONS OF PARTICULATE ORGANIC CARBON IN THE MEDITERRANEAN SEA AND THEIR RELEVANCE TO THE DEEP-LIVING COCCOLITHOPHORID CYCLOCOCCOLITHUS FRAGILIS. In Limnology and Oceanography (Vol. 13, Issue 4, pp. 693–697). John Wiley & Sons, Ltd. https://doi.org/10.4319/lo.1968.13.4.0693

Frankel, E. B., Shankar, R., Moresco, J. J., Yates, J. R., Volkmann, N., & Audhya, A. (2017). Ist1 regulates ESCRT-III assembly and function during multivesicular endosome biogenesis in Caenorhabditis elegans embryos. Nature Communications. https://doi.org/10.1038/s41467-017-01636-8

Friebele, E. S., Correll, D. L., & Faust, M. A. (1978). Relationship between phytoplankton cell size and the rate of orthophosphate uptake: in situ observations of an estuarine population. Marine Biology, 45(1), 39–52. https://doi.org/10.1007/BF00388976

Gemmell, B. J., Oh, G., Buskey, E. J., & Villareal, T. A. (2016). Dynamic sinking behaviour in marine phytoplankton: Rapid changes in buoyancy may aid in nutrient uptake. Proceedings of the Royal Society B: Biological Sciences, 283(1840). https://doi.org/10.1098/rspb.2016.1126

Gillmor, C. S., Roeder, A. H. K., Sieber, P., Somerville, C., & Lukowitz, W. (2016). A Genetic Screen for Mutations Affecting Cell Division in the Arabidopsis thaliana Embryo Identifies Seven Loci Required for Cytokinesis. PLOS ONE, 11(1), e0146492. https://doi.org/10.1371/JOURNAL.PONE.0146492

Glockling, S. L., Marshall, W. L., & Gleason, F. H. (2013). Phylogenetic interpretations and ecological potentials of the Mesomycetozoea (Ichthyosporea). In Fungal Ecology (Vol. 6, Issue 4, pp. 237–247). Elsevier Ltd. https://doi.org/10.1016/j.funeco.2013.03.005

Grau-Bové, X., Torruella, G., Donachie, S., Suga, H., Leonard, G., Richards, T. A., & Ruiz-Trillo, I. (2017). Dynamics of genomic innovation in the unicellular ancestry of animals. ELife. https://doi.org/10.7554/eLife.26036

Grosberg, R. K., & Strathmann, R. R. (2007). The evolution of multicellularity: A minor major transition? In Annual Review of Ecology, Evolution, and Systematics. https://doi.org/10.1146/annurev.ecolsys.36.102403.114735

Hadjebi, O., Casas-Terradellas, E., Garcia-Gonzalo, F. R., & Rosa, J. L. (2008). The RCC1 superfamily: From genes, to function, to disease. In Biochimica et Biophysica Acta - Molecular Cell Research. https://doi.org/10.1016/j.bbamcr.2008.03.015

Hassett, B. T., López, J. A., & Gradinger, R. (2015). Two New Species of Marine Saprotrophic Sphaeroformids in the Mesomycetozoea Isolated From the Sub-Arctic Bering Sea. Protist. https://doi.org/10.1016/j.protis.2015.04.004

Herron, M. D., Borin, J. M., Boswell, J. C., Walker, J., Chen, I. C. K., Knox, C. A., Boyd, M., Rosenzweig, F., & Ratcliff, W. C. (2019). De novo origins of multicellularity in response to predation. Scientific Reports. https://doi.org/10.1038/s41598-019-39558-8

Hirono, M., & Yoda, A. (1997). Isolation and Phenotypic Characterization of ChlamydomonasMutants Defective in Cytokinesis. CELL STRUCTURE AND FUNCTION, 22, 1–5.

Huang, H., Fletcher, L., Beeharry, N., Daniel, R., Kao, G., Yen, T. J., & Muschel, R. J. (2008). Abnormal Cytokinesis after X-Irradiation in Tumor Cells that Override the G2 DNA Damage Checkpoint. Cancer Research, 68(10), 3724–3732. https://doi.org/10.1158/0008-5472.CAN-08-0479

Hübner, S., Bahr, C., Gößmann, H., Efthymiadis, A., & Drenckhahn, D. (2003). Mitochondrial and nuclear localization of kanadaptin. European Journal of Cell Biology. https://doi.org/10.1078/0171-9335-00308

Hübner, S., Jans, D. A., Xiao, C. Y., John, A. P., & Drenckhahn, D. (2002). Signal- and importin-dependent nuclear targeting of the kidney anion exchanger 1-binding protein kanadaptin. Biochemical Journal. https://doi.org/10.1042/0264-6021:3610287

Jøstensen, J. P., Sperstad, S., Johansen, S., & Landfald, B. (2002). Molecular-phylogenetic, structural and biochemical features of a cold-adapted, marine ichthyosporean near the animal-fungal divergence, described from in vitro cultures. European Journal of Protistology. https://doi.org/10.1078/0932-4739-00855

Kadowaki, T., Zhao, Y., & Tartakoff, A. M. (1992). A conditional yeast mutant deficient in mRNA transport from nucleus to cytoplasm. Proceedings of the National Academy of Sciences, 89(6), 2312–2316. https://doi.org/10.1073/PNAS.89.6.2312

Kawecki, T. J., Lenski, R. E., Ebert, D., Hollis, B., Olivieri, I., & Whitlock, M. C. (2012). Experimental evolution. In Trends in Ecology and Evolution. https://doi.org/10.1016/j.tree.2012.06.001

Knoll, A. H. (2011). The Multiple Origins of Complex Multicellularity. Annual Review of Earth and Planetary Sciences, 39(1), 217–239. https://doi.org/10.1146/annurev.earth.031208.100209

Konopka, C. A., Schleede, J. B., Skop, A. R., & Bednarek, S. Y. (2006). Dynamin and cytokinesis. In Traffic. https://doi.org/10.1111/j.1600-0854.2006.00385.x

Koschwanez, J. H., Foster, K. R., & Murray, A. W. (2013). Improved use of a public good selects for the evolution of undifferentiated multicellularity. ELife, 2013(2). https://doi.org/10.7554/eLife.00367

Leigh, E. G., Smith, J. M., & Szathmary, E. (1995). The Major Transitions of Evolution. Evolution. https://doi.org/10.2307/2410462

Levin, T. C., Greaney, A. J., Wetzel, L., & King, N. (2014). The Rosetteless gene controls development in the choanoflagellate S. rosetta. ELife. https://doi.org/10.7554/eLife.04070

M, A., MW, C., U, V., & J, A. (1990). A yeast mutant, PRP20, altered in mRNA metabolism and maintenance of the nuclear structure, is defective in a gene homologous to the human gene RCC1 which is involved in the control of chromosome condensation. Molecular & General Genetics : MGG, 224(1), 72–80. https://doi.org/10.1007/BF00259453

M, O., H, O., & T, N. (1989). The RCC1 protein, a regulator for the onset of chromosome condensation locates in the nucleus and binds to DNA. The Journal of Cell Biology, 109(4 Pt 1), 1389–1397. https://doi.org/10.1083/JCB.109.4.1389

Marshall, N. B. (1954). *Aspects of deep sea biology* (380 pp). Hutchinson.

Marshall, W. L., & Berbee, M. L. (2013). Comparative morphology and genealogical delimitation of cryptic species of sympatric isolates of sphaeroforma (Ichthyosporea, Opisthokonta). Protist. https://doi.org/10.1016/j.protis.2012.12.002

Marshall, W. L., Celio, G., McLaughlin, D. J., & Berbee, M. L. (2008). Multiple Isolations of a Culturable, Motile Ichthyosporean (Mesomycetozoa, Opisthokonta), Creolimax fragrantissima n. gen., n. sp., from Marine Invertebrate Digestive Tracts. Protist. https://doi.org/10.1016/j.protis.2008.03.003

Masud Rana, A. Y. K. M., Tsujioka, M., Miyagishima, S., Ueda, M., & Yumura, S. (2013). Dynamin contributes to cytokinesis by stabilizing actin filaments in the contractile ring. Genes to Cells. https://doi.org/10.1111/gtc.12060

Mendoza, L., Taylor, J. W., & Ajello, L. (2002). The class Mesomycetozoea: A heterogeneous group of microorganisms at the animal-fungal boundary. In Annual Review of Microbiology (Vol. 56, pp. 315–344). Annu Rev Microbiol. https://doi.org/10.1146/annurev.micro.56.012302.160950

Millero, F. J., & Huang, F. (2009). The density of seawater as a function of salinity (5 to 70 g kg -1) and temperature (273.15 to 363.15 K). Ocean Science, 5(2), 91–100. https://doi.org/10.5194/os-5-91-2009

Nanninga, N. (2001). Cytokinesis in Prokaryotes and Eukaryotes: Common Principles and Different Solutions. Microbiology and Molecular Biology Reviews, 65(2), 319. https://doi.org/10.1128/MMBR.65.2.319-333.2001

Niklas, K. J., & Newman, S. A. (2013). The origins of multicellular organisms. Evolution and Development, 15(1), 41–52. https://doi.org/10.1111/ede.12013

Ondracka, A., Dudin, O., & Ruiz-Trillo, I. (2018). Decoupling of Nuclear Division Cycles and Cell Size during the Coenocytic Growth of the Ichthyosporean Sphaeroforma arctica. Current Biology, 28(12). https://doi.org/10.1016/j.cub.2018.04.074

Oud, B., Guadalupe-Medina, V., Nijkamp, J. F., De Ridder, D., Pronk, J. T., Van Maris, A. J. A., & Daran, J. M. (2013). Genome duplication and mutations in ACE2 cause multicellular, fast-sedimenting phenotypes in evolved Saccharomyces cerevisiae. Proceedings of the National Academy of Sciences of the United States of America, 110(45), E4223–E4231. https://doi.org/10.1073/pnas.1305949110

Parfrey, L. W., & Lahr, D. J. G. (2013). Multicellularity arose several times in the evolution of eukaryotes. BioEssays. https://doi.org/10.1002/bies.201200143

Parra-Acero, H., Harcet, M., Sánchez-Pons, N., Casacuberta, E., Brown, N. H., Dudin, O., & Ruiz-Trillo, I. (2020). Integrin-Mediated Adhesion in the Unicellular Holozoan Capsaspora owczarzaki. Current Biology. https://doi.org/10.1016/j.cub.2020.08.015

Pérez-Posada, A., Dudin, O., Ocaña-Pallarès, E., Ruiz-Trillo, I., & Ondracka, A. (2020). Cell cycle transcriptomics of Capsaspora provides insights into the evolution of cyclin-CDK machinery. PLoS Genetics, 16(3). https://doi.org/10.1371/journal.pgen.1008584

Pfeifer, F. (2015). Haloarchaea and the formation of gas vesicles. In Life. https://doi.org/10.3390/life5010385

Prabhu, A. V., Luu, W., Sharpe, L. J., & Brown, A. J. (2016). Cholesterol-mediated degradation of 7-dehydrocholesterol reductase switches the balance from cholesterol to Vitamin D synthesis. Journal of Biological Chemistry. https://doi.org/10.1074/jbc.M115.699546

Qiao, L., Zheng, J., Tian, Y., Zhang, Q., Wang, X., Chen, J. J., & Zhang, W. (2018). Regulator of chromatin condensation 1 abrogates the G1 cell cycle checkpoint via Cdk1 in human papillomavirus E7-expressing epithelium and cervical cancer cells article. Cell Death and Disease. https://doi.org/10.1038/s41419-018-0584-z

Queller, D. C., Ponte, E., Bozzaro, S., & Strassmann, J. E. (2003). Single-gene greenbeard effects in the social amoeba Dictyostelium discoideum. Science. https://doi.org/10.1126/science.1077742

Ratcliff, W. C., Denison, R. F., Borrello, M., & Travisano, M. (2012). Experimental evolution of multicellularity. Proceedings of the National Academy of Sciences of the United States of America. https://doi.org/10.1073/pnas.1115323109

Ratcliff, W. C., Fankhauser, J. D., Rogers, D. W., Greig, D., & Travisano, M. (2015). Origins of multicellular evolvability in snowflake yeast. Nature Communications. https://doi.org/10.1038/ncomms7102

Ratcliff, W. C., Herron, M. D., Howell, K., Pentz, J. T., Rosenzweig, F., & Travisano, M. (2013). Experimental evolution of an alternating uni- and multicellular life cycle in Chlamydomonas reinhardtii. Nature Communications. https://doi.org/10.1038/ncomms3742

Rikhy, R., Mavrakis, M., & Lippincott-Schwartz, J. (2015). Dynamin regulates metaphase furrow formation and plasma membrane compartmentalization in the syncytial Drosophila embryo. Biology Open. https://doi.org/10.1242/bio.20149936

Rohlfs, M., Arasada, R., Batsios, P., Janzen, J., & Schleicher, M. (2007). The Ste20-like kinase SvkA of Dictyostelium discoideum is essential for late stages of cytokinesis. Journal of Cell Science. https://doi.org/10.1242/jcs.012179

Rokas, A. (2008). The molecular origins of multicellular transitions. In Current Opinion in Genetics and Development (Vol. 18, Issue 6, pp. 472–478). Elsevier Current Trends. https://doi.org/10.1016/j.gde.2008.09.004

Ruiz-Trillo, I., Burger, G., Holland, P. W. H., King, N., Lang, B. F., Roger, A. J., & Gray, M. W. (2007). The origins of multicellularity: a multi-taxon genome initiative. Trends in Genetics, 23(3), 113–118. https://doi.org/10.1016/j.tig.2007.01.005

Ruiz-Trillo, I., & de Mendoza, A. (2020). Towards understanding the origin of animal development. Development (Cambridge, England). https://doi.org/10.1242/dev.192575

Sebé-Pedrós, A., Degnan, B. M., & Ruiz-Trillo, I. (2017). The origin of Metazoa: A unicellular perspective. In Nature Reviews Genetics. https://doi.org/10.1038/nrg.2017.21

Sebé-Pedrós, A., Irimia, M., del Campo, J., Parra-Acero, H., Russ, C., Nusbaum, C., Blencowe, B. J., & Ruiz-Trillo, I. (2013). Regulated aggregative multicellularity in a close unicellular relative of metazoa. ELife, 2(2:e01287). https://doi.org/10.7554/eLife.01287

Smayda, & J., T. (1970). The suspension and sinking of phytoplankton in the sea. Oceanogr. Mar. Biol. Ann. Rev., 8, 353–414. https://ci.nii.ac.jp/naid/10009425107

Smayda, T. J. (1971). Normal and accelerated sinking of phytoplankton in the sea. Marine Geology, 11(2), 105–122. https://doi.org/10.1016/0025-3227(71)90070-3

Smit, A., Hubley, R., & Grenn, P. (2015). RepeatMasker Open-4.0. RepeatMasker Open.

Strand, E., Jørgensen, C., & Huse, G. (2005). Modelling buoyancy regulation in fishes with swimbladders: Bioenergetics and behaviour. Ecological Modelling. https://doi.org/10.1016/j.ecolmodel.2004.12.013

Strassmann, J. E., Gilbert, O. M., & Queller, D. C. (2011). Kin discrimination and cooperation in microbes. In Annual Review of Microbiology. https://doi.org/10.1146/annurev.micro.112408.134109

Suga, H., & Ruiz-Trillo, I. (2013). Development of ichthyosporeans sheds light on the origin of metazoan multicellularity. Developmental Biology. https://doi.org/10.1016/j.ydbio.2013.01.009

Sundby, S., & Kristiansen, T. (2015). The principles of buoyancy in Marine Fish Eggs and their vertical distributions across the World Oceans. PLoS ONE. https://doi.org/10.1371/journal.pone.0138821

Tarnita, C. E., Taubes, C. H., & Nowak, M. A. (2013). Evolutionary construction by staying together and coming together. Journal of Theoretical Biology. https://doi.org/10.1016/j.jtbi.2012.11.022

Villareal, T. A., & Carpenter, E. J. (2003). Buoyancy regulation and the potential for vertical migration in the oceanic cyanobacterium Trichodesmium. Microbial Ecology. https://doi.org/10.1007/s00248-002-1012-5

Waite, A., Fisher, A., Thompson, P., & Harrison, P. (1997). Sinking rate versus cell volume relationships illuminate sinking rate control mechanisms in marine diatoms. Marine Ecology Progress Series, 157, 97–108. https://doi.org/10.3354/meps157097

Wang, M. C., Lu, Y., & Baldock, C. (2009). Fibrillin Microfibrils: A Key Role for the Interbead Region in Elasticity. Journal of Molecular Biology. https://doi.org/10.1016/j.jmb.2009.02.062

Wang, Y., Wang, R., & Tang, D. D. (2020). Ste20-like kinase-mediated control of actin polymerization is a new mechanism for thin filament-associated regulation of airway smooth muscle contraction. American Journal of Respiratory Cell and Molecular Biology. https://doi.org/10.1165/RCMB.2019-0310OC

Wickham, H. (2016). ggplot2: Elegant Graphics for Data Analysis. In Journal of the Royal Statistical Society: Series A (Statistics in Society).

Wielgoss, S., Wolfensberger, R., Sun, L., Fiegna, F., & Velicer, G. J. (2019). Social genes are selection hotspots in kin groups of a soil microbe. Science. https://doi.org/10.1126/science.aar4416

Xiao, J., Chen, X. W., Davies, B. A., Saltiel, A. R., Katzmann, D. J., & Xu, Z. (2009). Structural basis of Ist1 function and Ist1-Did2 interaction in the multivesicular body pathway and cytokinesis. Molecular Biology of the Cell. https://doi.org/10.1091/mbc.E09-05-0403

Yin, W., Kim, H. T., Wang, S. P., Gunawan, F., Li, R., Buettner, C., Grohmann, B., Sengle, G., Sinner, D., Offermanns, S., & Stainier, D. Y. R. (2019). Fibrillin-2 is a key mediator of smooth muscle extracellular matrix homeostasis during mouse tracheal tubulogenesis. European Respiratory Journal. https://doi.org/10.1183/13993003.00840-2018

Zhang, H., Apfelroth, S. D., Hu, W., Davis, E. C., Sanguineti, C., Bonadio, J., Mecham, R. P., & Ramirez, F. (1994). Structure and expression of fibrillin-2, a novel microfibrillar component preferentially located in elastic matrices. Journal of Cell Biology. https://doi.org/10.1083/jcb.124.5.855

