## Supplementary material for "Regulation of sedimentation rate shapes the evolution of multicellularity in a unicellular relative of animals": Figure 2 - Source Data 1

**Figure 2- Source Data 1. Inference of doubling times and effective population size [Ne] from direct estimates of cell numbers and nuclei per cells over the time course of two transfers.**

| Timepoint [h] | Cell numbers[1] | Nuclei (per cell)[1] | Nuclei (total)[2] | Nucleic doublings[3,4] | Population size [Ne][5] |
|---|---|---|---|---|---|
| 0 | 1.65E+04 | 1.82 | 3.00E+04 | | |
| 24 | 1.90E+04 | 9.69 | 1.84E+05 | **2.62** | **7.86 E+04** |
| 24 | 6.00E+02 | 9.69 | 5.81E+03 | | |
| 48 | 6.00E+05 | 14.8 | 8.84E+06 | **10.6** | **6.14E+04** |

[1] determined experimentally

[2] calculated as product of *cell number* and *nuclei (per cell)*

[3] value at 24h: calculated as LOG2 (Nuclei,total) [24h] – LOG2 (Nuclei,total) [0h]

[4] value at 48h: calculated as LOG2 (Nuclei,total) [48h] – LOG2 (Nuclei,total) [24h]

[5] calculated as product of start (=bottleneck) cell numbers and nucleic doublings
