## Supplementary material for "Regulation of sedimentation rate shapes the evolution of multicellularity in a unicellular relative of animals": Figure 4 - Source Data 1

### Figure 4- Source Data 1. Genome resequencing summary: data processing and mapping.

| ID | Treatment (*) | Clumping Type (**) | Total Reads (***) | Raw Data (Gb) | Trimmed Reads (%) (****) | TrimmedData (Gb) | Aligned Data (%) (*****) | Deduplicated Data (%) | Informative Data (Gb) | Fraction of Reference Genome covered (%) | Effective Average Coverage |
|---|---|---|---|---|---|---|---|---|---|---|---|
| AN | A | A | 64,961,014 | 8.12 | 44.6 | 4.50 | 99.31 | 96.74 | 4.35 | 88 | 34.5 |
| S1 | E | H | 66,575,790 | 8.32 | 39.9 | 5.00 | 99.15 | 97.28 | 4.86 | 87 | 38.9 |
| S2 | E | I | 60,999,004 | 7.62 | 40.5 | 4.54 | 99.25 | 97.46 | 4.42 | 87 | 35.4 |
| S3 | E | I | 59,917,632 | 7.49 | 39.6 | 4.52 | 99.14 | 97.14 | 4.39 | 87 | 35.1 |
| S4 | E | H | 57,928,616 | 7.24 | 40.5 | 4.31 | 99.24 | 97.35 | 4.19 | 87 | 33.8 |
| S5 | E | I | 57,822,932 | 7.23 | 41.7 | 4.21 | 99.28 | 97.42 | 4.10 | 87 | 32.9 |
| S6 | E | L | 57,657,624 | 7.21 | 40.0 | 4.32 | 99.29 | 97.50 | 4.22 | 88 | 33.6 |
| S7 | E | I | 52,765,444 | 6.60 | 39.2 | 4.01 | 99.27 | 96.97 | 3.89 | 88 | 30.7 |
| S8 | E | L | 52,155,682 | 6.52 | 42.2 | 3.77 | 99.35 | 97.16 | 3.66 | 87 | 29.3 |
| S9 | E | H | 59,011,232 | 7.38 | 39.9 | 4.43 | 99.24 | 97.24 | 4.31 | 89 | 33.9 |
| S10 | E | L | 57,388,476 | 7.17 | 40.1 | 4.30 | 99.47 | 96.84 | 4.16 | 90 | 32.2 |
| Total | | | 58,834,859 | 7.35 | 40.8 | 4.36 | 99.27 | 97.19 | 4.23 | 88 | 33.7 |
| Control: | | | | | | | | | | | |
| DRR183670 | R | C | 116,162,364 | 29.0 | 45.4 | 15.9 | 98.37 | 96.63 | 15.3 | 94 | 114 |

(*) A, ancestral; E, evolved; R, reference
(**) A, ancestral; H, highly clumpy; I, Intermediate clumpiness; L, Low clumpiness; C, control
(***) Uniform raw read length of 125bp
(****) Data filtered during trimming prior to mapping
(*****) Given a reference genome size of 142,721,209bp
