## Supplementary material for "Regulation of sedimentation rate shapes the evolution of multicellularity in a unicellular relative of animals": Figure 4 - Source Data 2

### Figure 4- Source Data 2. Summary and classification of annotated repeat regions in the *Sphaeroforma arctica* reference genome.

| Name and classification of element | Number of elements | Length occupied [bp] | Percentage of total sequence[1] [%] |
|---|---|---|---|
| **Retroelements** | **44,831** | **5,606,241** | **3.99** |
| SINEs: | 302 | 24,043 | 0.02 |
| Penelope | 3,232 | 256,219 | 0.18 |
| LINEs: | 29,324 | 3,997,048 | 2.84 |
| CRE/SLACS | 22 | 1078 | 0 |
| L2/CR1/Rex | 22,777 | 3,502,253 | 2.49 |
| R1/LOA/Jockey | 375 | 27,235 | 0.02 |
| R2/R4/NeSL | 126 | 8,005 | 0.01 |
| RTE/Bov-B | 479 | 38,312 | 0.03 |
| L1/CIN4 | 1,779 | 117,738 | 0.08 |
| LTR elements: | 15,205 | 1,585,150 | 1.13 |
| BEL/Pao | 469 | 32,962 | 0.02 |
| Ty1/Copia | 651 | 145,765 | 0.1 |
| Gypsy/DIRS1 | 7,466 | 999,654 | 0.71 |
| Retroviral | 5,423 | 322,159 | 0.23 |
| **DNA transposons** | **89,667** | **8,369,254** | **5.95** |
| hobo-Activator | 8,125 | 472,782 | 0.34 |
| Tc1-IS630-Pogo | 12,356 | 1,604,248 | 1.14 |
| En-Spm | 0 | 0 | 0 |
| MuDR-IS905 | 0 | 0 | 0 |
| PiggyBac | 125 | 6,753 | 0 |
| Tourist/Harbinger | 1,314 | 67,309 | 0.05 |
| Other: Mirage, P-element, Transib | 2,894 | 195,557 | 0.14 |
| **Rolling-circles** | **4,725** | **574,140** | **0.41** |
| **Unclassified** | **389** | **51,677** | **0.04** |
| **Total interspersed repeats** | **-** | **14,027,172** | **9.97** |
| **Small RNAs** | **1,146** | **1,193,846** | **0.85** |
| **Satellites** | **1,861** | **345,389** | **0.25** |
| **Simple repeats** | **73,308** | **12,722,055** | **9.05** |
| **Low complexity** | **2,982** | **166,665** | **0.12** |

[1] Total reference genome size (Sarc4) is 142,721,209 bp distributed across 1,750 contigs with a GC-content of ~38.5%.
Masking of the entire portion (~20.6%) of annotated repetitive sequence resulted in a final repeat-masked genome length of 113,693,024 bp.
Beware that the lengths and percentage of total sequence do not directly add up to yield the above numbers, since repetitive regions may house overlapping annotations for different elements (e.g., simple repeats within other elements).
