## Supplementary material for "Regulation of sedimentation rate shapes the evolution of multicellularity in a unicellular relative of animals": Figure 4 - Source Data 3

### Figure 4- Source Data 3. Variant calling

| | | | | | | | | **FAST** | | | **MEDIUM** | | | | **SLOW** | | | |
|---|---|---|---|---|---|---|---|---|---|---|---|---|---|---|---|---|---|---|
| ID | Seq id | Position | TYPE | Effect | Mutation | Annotation[1] | Affected genes | **S01** | **S04** | **S09** | **S02** | **S03** | **S05** | **S07** | **S06** | **S08** | **S10** | Read support[2] |
| 1 | scaffold658 | 3,101 | SNP | SY | C→T | T89T (AC**C**-AC**T**) | Sarc4_g12019T | **1** | | | | | | | | | | 17/17 |
| 2 | scaffold052 | 23,219 | SNP | NS | A→G | E90G (G**A**A→G**G**A) | Sarc4_g23215T | | **1** | | | | | | | | | 25/25 |
| 3 | scaffold209 | 85,868 | SNP | NC | G→C | intronic **(-314/+740**) | Sarc4_g32431T | | **1** | | | | | | | | | 29/29 |
| 4 | scaffold224 | 61,451 | SNP | NS | T→A | S102C (**A**GC→**T**GC) | Sarc4_g33124T | | **1** | | | | | | | | | 12/12 |
| 5 | scaffold226 | 196,494 | SNP | IN | G→A | intergenic **(-587**/+1084) | Sarc4_g33270T | | **1** | | | | | | | | | 27/27 |
| 6 | scaffold617 | 26,290 | SNP | NC | C→T | intronic **(+390/-279**) | Sarc4_g11520T | | **1** | | | | | | | | | 30/30 |
| 7 | scaffold042 | 318,312 | SNP | NC | C→T | intronic **(+102/-711**) | Sarc4_g22580T | | | **1** | | | | | | | | 32/32 |
| 8 | scaffold278 | 57,886 | SNP | NC | A→G | intronic **(+291/-424**) | Sarc4_g3019T | | | **1** | | | | | | | | 33/33 |
| 9 | scaffold426 | 46,771 | SNP | NC | A→G | intronic **(+280/-543**) | Sarc4_g7653T | | | **1** | | | | | | | | 21/21 |
| 10 | scaffold645 | 9,116 | SNP | NS | C→T | A923V (G**C**A→G**T**A) | Sarc4_g11880T | | | **1** | | | | | | | | 22/22 |
| 11 | scaffold651 | 57,503 | SNP | IN | A→T | intergenic **(+186/+1338**) | Sarc4_g11957 / Sarc4_g11958 | | | **1** | | | | | | | | 30/30 |
| 12 | scaffold241 | 79,469 | SNP | IN | G→A | intergenic **(-268**/+232) | Sarc4_g33887T | | | | **1** | | | | | | | 23/23 |
| 13 | scaffold406 | 53,611 | SNP | NS | C→G | V176L (**G**TT→**C**TT) | Sarc4_g7170T | | | | **1** | | | | | | | 29/29 |
| 14 | scaffold167 | 184,318 | INS | FS | +AG | coding (**85**/117 nt) | Sarc4_g30293T | | | | | **1** | | | | | | 12/12 |
| 15 | scaffold381 | 83,481 | SNP | NS | C→T | G148S (**G**GT→**A**GT) | Sarc4_g6394T | | | | | **1** | | | | | | 24/26 |
| 16 | scaffold153 | 151,457 | SNP | IN | G→T | intergenic (+512**/-553**) | Sarc4_g29511T | | | | | | **1** | | | | | 35/35 |
| 17 | scaffold414 | 97,263 | SNP | NS | C→T | Q1750* (**C**AG→**T**AG) | Sarc4_g7365T | | | | | | **1** | | | | | 23/23 |
| 18 | scaffold731 | 5,386 | SNP | NC | A→G | intronic (–/+118) | Sarc4_g12953 | | | | | | **1** | | | | | 28/28 |
| 19 | scaffold301 | 38,194 | SNP | NS | C→T | D25N (**G**AC→**A**AC) | Sarc4_g3900T | **1** | | | | | | **1** | | | | 37/37; 27/27 |
| 20 | scaffold193 | 291,552 | INS | IN | +G | intergenic (+3610**/-4749**) | Sarc4_g31776T | | | | | | | | **1** | | | 27/27 |
| 21 | scaffold209 | 35,723 | SNP | NC | C→T | intronic **(+360/-796**) | Sarc4_g32417T | | | | | | | | **1** | | | 28/28 |
| 22 | scaffold336 | 12,377 | SNP | NC | G→A | intronic **(+3872/-76**) | Sarc4_g4950T | | | | | | | | **1** | | | 25/25 |
| 23 | scaffold842 | 3,222 | SNP | IN | C→T | intergenic (**–**/+1674) | Sarc4_g14312 | | | | | | | | | **1** | | 18/21 |
| 24 | scaffold021 | 360,117 | SNP | SY | T→C | P45P (CC**T**→CC**C**) | Sarc4_g18855 | | | | | | | | | | **1** | 18/18 |
| 25 | scaffold207 | 143,138 | SNP | NS | G→A | R284C (**C**GC→**T**GC) | Sarc4_g32374T | | | | | | | | | | **1** | 17/17 |

SNP, single nucleotide polymorphism; INS, Insertion; NS, non-synonymous (including non-sense) mutations; SY, synonymous mutations; NC, non-coding (i.e., intronic) mutations; IN, intergenic mutations; FS, frame-shift mutation

[1] according to breseq's gdtools ANNOTATE (**bold**: changed base in codon (for NS mutations); position down/upstream of previous/next exon (for NC mutations); position down/upstream of previous/next gene (for IN mutations); position of insertion in coding sequence (for FS mutations))

[2] required variant allele support: x ≥ 10, required majority frequency ≥ 0.8.
