## Supplementary material for "Regulation of sedimentation rate shapes the evolution of multicellularity in a unicellular relative of animals": Figure 4 - Source Data 4

### Figure 4- Source Data 4. Transcript abundance of affected genes during *S. arctica* native life-cycle

| ID | Affected genes | TYPE | Effect | Transcript abundance (in tpm)[1] | | | | | | | | | |
|---|---|---|---|---|---|---|---|---|---|---|---|---|---|
| | | | | 12h | 18h | 24h | 30h | 36h | 42h | 48h | 54h | 60h | 66h |
| 1 | Sarc4_g12019T | SNP | SY | 34.3687 | 37.1965 | 35.0699 | 37.0085 | 44.8557 | 97.1095 | 98.9791 | 86.1972 | 83.1505 | 61.6903 |
| 2 | Sarc4_g23215T | SNP | NS | 19.6851 | 19.1455 | 17.5439 | 16.2834 | 23.1461 | 18.6644 | 17.9979 | 19.4054 | 17.2579 | 11.882 |
| 3 | Sarc4_g32431T | SNP | NC | 69.5583 | 76.5122 | 76.5697 | 74.6316 | 91.2736 | 100.101 | 106.021 | 124.43 | 161.934 | 160.266 |
| 4 | Sarc4_g33124T | SNP | NS | 0 | 0 | 0 | 0 | 0 | 0 | 0 | 0 | 0 | 0 |
| 5 | Sarc4_g33270T | SNP | IN | 22.9663 | 28.5196 | 22.2241 | 23.1924 | 27.1365 | 21.835 | 19.1085 | 15.252 | 13.0743 | 11.0699 |
| 6 | Sarc4_g11520T | SNP | NC | 79.2703 | 83.0262 | 83.5683 | 75.0504 | 86.8678 | 124.211 | 171.462 | 169.185 | 127.728 | 72.5625 |
| 7 | Sarc4_g22580T | SNP | NC | 1.29071 | 1.55378 | 0.82560 | 0.84584 | 0.8672 | 1.17345 | 1.70523 | 2.04122 | 3.36911 | 4.0704 |
| 8 | Sarc4_g3019T | SNP | NC | 114.017 | 108.556 | 110.86 | 101.407 | 122.306 | 120.344 | 116.719 | 110.999 | 101.797 | 88.9507 |
| 9 | Sarc4_g7653T | SNP | NC | 16.3923 | 17.5438 | 16.7544 | 18.797 | 20.5921 | 19.6488 | 22.9749 | 23.9413 | 19.1717 | 15.4509 |
| 10 | Sarc4_g11880T | SNP | NS | 6.91487 | 6.60516 | 4.32566 | 4.51767 | 6.66984 | 4.26517 | 3.17804 | 2.43333 | 1.54733 | 2.50654 |
| 11 | Sarc4_g11957 | SNP | IN | 0.55330 | 0.55777 | 0 | 0 | 0.33447 | 0.39681 | 0.84829 | 0 | 0 | 0 |
| | Sarc4_g11958 | | | 0 | 0 | 0 | 0 | 0 | 0 | 0 | 0 | 0 | 0 |
| 12 | Sarc4_g33887T | SNP | IN | 0.25507 | 1.01635 | 0 | 0 | 0 | 0 | 0 | 0 | 0 | 0 |
| 13 | Sarc4_g7170T | SNP | NS | 0.31083 | 0.10386 | 0.11287 | 0.21621 | 0 | 0.37790 | 0 | 0.12197 | 0 | 0.28910 |
| 14 | Sarc4_g30293T | INS | FS | 23.7182 | 29.8256 | 31.1292 | 41.2785 | 57.8007 | 64.8248 | 71.5185 | 70.6809 | 60.6227 | 54.8389 |
| 15 | Sarc4_g6394T | SNP | NS | 19.3595 | 22.3913 | 19.4656 | 14.576 | 17.1674 | 28.6616 | 43.6442 | 131.04 | 64.7505 | 18.36 |
| 16 | Sarc4_g29511T | SNP | IN | 0 | 0.14618 | 0 | 0 | 0.15940 | 0 | 0.49779 | 0.17462 | 0 | 0.17259 |
| 17 | Sarc4_g7365T | SNP | NS | 41.1783 | 31.772 | 22.8192 | 27.044 | 34.9059 | 28.5977 | 19.6541 | 11.4106 | 12.3447 | 32.656 |
| 18 | Sarc4_g12953 | SNP | NC | 57.216 | 69.811 | 67.439 | 66.957 | 97.928 | 115.4 | 118.91 | 129.18 | 142.42 | 131.73 |
| 19 | Sarc4_g3900T | SNP | NS | 0 | 0 | 0 | 0 | 0 | 0 | 0 | 0 | 0 | 0 |
| 20 | Sarc4_g31776T | INS | IN | 0 | 0 | 0 | 0 | 0 | 0 | 0 | 0 | 0 | 0 |
| 21 | Sarc4_g32417T | SNP | NC | 5.6145 | 6.5909 | 6.5687 | 7.5284 | 10.521 | 37.328 | 55.623 | 71.687 | 57.300 | 77.678 |
| 22 | Sarc4_g4950T | SNP | NC | 2.9573 | 2.0139 | 1.1597 | 2.5930 | 2.1831 | 7.3580 | 9.1223 | 12.325 | 18.977 | 22.469 |
| 23 | Sarc4_g14312 | SNP | IN | 0 | 0 | 0 | 0 | 0 | 0 | 0 | 0 | 0.1011 | 0.1176 |
| 24 | Sarc4_g18855 | SNP | SY | 14.359 | 38.709 | 33.958 | 49.115 | 51.133 | 63.816 | 65.745 | 82.792 | 92.719 | 71.047 |
| 25 | Sarc4_g32374T | SNP | NS | 0 | 0.0286 | 0 | 0 | 0 | 0 | 0 | 0 | 0 | 0 |

[1] according to Dudin et al. 2019
