## Supplementary material for "Regulation of sedimentation rate shapes the evolution of multicellularity in a unicellular relative of animals": Figure 4 - Source Data 5

### Figure 4- Source Data 5. Annotation and gene prediction of mutational targets.

| ID | Locus tag | **Gene name[1]** | **Annotation[1]** | **COG assignment[1]** |
|---|---|---|---|---|
| 1 | Sarc4_g12019T | Ist1 | Regulator of Vps4 activity in the multivesicular bodies (MVB) pathway | - |
| 2 | Sarc4_g23215T | SLC4A1AP, Kanadaptin | Solute carrier family 4 (Anion exchanger), member 1, adaptor protein | Signal Transduction (T) |
| 3 | Sarc4_g32431T | SLK | ste20-like kinase adaptor | Signal Transduction (T) |
| 4 | Sarc4_g33124T | - | - | - |
| 5 | Sarc4_g33270T | - | - | Function Unknown (S) |
| 6 | Sarc4_g11520T | DHCR7 | delta(14)-sterol reductase, ERG4_ERG24 | Lipid metabolism (I) |
| 7 | Sarc4_g22580T | DUF563 | Domain of unknown function DUF563 | - |
| 8 | Sarc4_g3019T | RCC1 | Regulator of chromosome condensation (RCC1) repeat | Post-translational modification, protein turnover, chaperone functions (O) |
| 9 | Sarc4_g7653T | HLH | Helix-loop-helix DNA-binding, PHD finger superfamily; cl22851 | - |
| 10 | Sarc4_g11880T | C2H2 | Zn finger nucleotide-binding | - |
| 11 | Sarc4_g11957T / Sarc4_g11958T | - | - | - |
| 12 | Sarc4_g33887T | - | - | - |
| 13 | Sarc4_g7170T | - | - | - |
| 14 | Sarc4_g30293T | DAO | D-amino-acid oxidase | Intracellular trafficking and secretion (U) |
| 15 | Sarc4_g6394T | Erv26 | Transmembrane adaptor | Lipid metabolism (I), Signal Transduction (T) |
| 16 | Sarc4_g29511T | - | - | - |
| 17 | Sarc4_g7365T | Fibrillin-2 | Lectin C-type domain, EGF repeat-like domain | Signal Transduction (T) |
| 18 | Sarc4_g12953T | DNM1 | Dynamin-related protein | Intracellular trafficking and secretion (U) |
| 19 | Sarc4_g3900T | - | - | - |
| 20 | Sarc4_g31776T | - | - | - |
| 21 | Sarc4_g32417T | IRK | Potassium inwardly-rectifying k+ (IRK) channel subfamily J, member (KCNJ6) | Inorganic ion transport and metabolism (P) |
| 22 | Sarc4_g4950T | - | - | - |
| 23 | Sarc4_g14312T | - | - | - |
| 24 | Sarc4_g18855T | RILP-like | Rab interacting lysosomal protein-like 1 and 2 | Function Unknown (S) |
| 25 | Sarc4_g32374T | RVT_2 | Retrotransposon protein | Function Unknown (S) |

**[1] adapted and extended from Grau-Bové et al. (2017) respectively**
